# Advanced Peptide Nanoparticles Enable Robust and Efficient delivery of gene editors across cell types

**DOI:** 10.1101/2024.11.27.624305

**Authors:** Oskar Gustafsson, Supriya Krishna, Sophia Borate, Marziyeh Ghaeidamini, Xiuming Liang, Osama Saher, Raul Cuellar, Björn K. Birdsong, Samantha Roudi, H. Yesid Estupiñán, Evren Alici, CI Edvard Smith, Elin K. Esbjörner, Simone Spuler, Olivier Gerrit de Jong, Helena Escobar, Joel Z. Nordin, Samir EL Andaloussi

## Abstract

Efficient delivery of the CRISPR/Cas9 system and its larger derivatives, base editors, and prime editors remain a significant challenge, particularly in tissue-specific stem cells and induced pluripotent stem cells (iPSCs). This study optimized a novel family of cell-penetrating peptides, hPep, to deliver gene-editing ribonucleoproteins. The hPep-based nanoparticles enable highly efficient and biocompatible delivery of Cre recombinase, Cas9, base-, and prime editors. Using base editors, robust and nearly complete genome editing was achieved in the human cells: HEK293T (96%), iPSCs (74%), and muscle stem cells (80%). This strategy opens promising avenues for ex vivo and, potentially, in vivo applications. Incorporating silica particles enhanced the system’s versatility, facilitating cargo-agnostic delivery. Notably, the nanoparticles can be synthesized quickly on a benchtop and stored as lyophilized powder without compromising functionality. This represents a significant advancement in the feasibility and scalability of gene-editing delivery technologies.

## Introduction

CRISPR/Cas9 and its derivatives, such as base editors (BEs) and prime editors (PEs), have emerged as the most powerful tools for genome editing. Compared to earlier technologies like Transcription activator-like effector nuclease and zinc finger nucleases, CRISPR offers simple targeting and modification of cellular DNA ^1,2^. However, the relatively large Cas9 protein and guide RNA (gRNA) present significant challenges for intracellular delivery, particularly when using adeno-associated viruses (AAVs) as delivery vectors. With their limited packaging capacity, AAVs often require the Cas9 coding sequence to be split into two vectors, complicating the delivery process. Additionally, viral delivery can result in prolonged Cas9 expression, raising concerns about on-and off-target genotoxicity ^3–7^.

Non-viral delivery methods, particularly lipid nanoparticle (LNP)-mediated Cas9 mRNA and gRNA delivery, have gained traction as an alternative. While LNPs offer promising solutions, delivering the gene-modifying enzyme as mRNA necessitates translation within the cell, and hence, such approaches heavily rely on maximizing gRNA stability to avoid its degradation during Cas9 synthesis. Moreover, mRNA delivery often leads to high levels of protein expression, potentially increasing off-target activity and immunogenicity ^8–11^.

Cas9-gRNA ribonucleoprotein (RNP) complex delivery confers significant advantages in gene editing due to its transient bioavailability, which minimizes off-target effects, eliminates the risk of genomic integration, and mitigates immune responses against the bacterially derived Cas9 protein ^12,13^. Despite these benefits, delivering Cas9 and its derivatives as RNP poses unique challenges. Unlike nucleic acids, Cas9 RNP lacks consistent charge and hydrophobicity, making it sensitive to denaturing buffers and complicating its effective delivery ^14^.

Thus, there is a pressing need for alternative methods to achieve efficient intracellular delivery of Cas9 RNP and its derivatives. Current strategies fall into two broad categories: physical and nanoparticle-based approaches. Physical methods, including microinjection, electroporation, and membrane deformation techniques, are highly effective but face limitations such as the need for specialized equipment, difficulty in scaling, and limited applicability for in vivo applications ^15–19^.

A promising alternative involves the use of cell-penetrating peptides (CPPs). These peptides can either be covalently attached to Cas9 or mixed to form nanoparticles with the Cas9 RNP, mainly by charge-based electrostatic interactions. While direct conjugation of CPPs with Cas9 creates a self-deliverable complex, this approach often requires large quantities of RNP due to inefficient intracellular delivery ^20–24^. The non-covalent strategy typically relies on charge interactions between the RNP and CPPs, leading to nanoparticle formation. This approach has proven efficient for cellular transfection and delivering various cargo types, such as RNPs and homology-directed repair (HDR) templates ^25^.

However, CPPs often suffer from poor serum stability, underscoring the critical need for scalable, serum-compatible delivery methods for Cas9 and its larger derivatives. Such systems are essential for gene-editing in cells and tissues, including iPSCs and skeletal muscle, where non-viral delivery remains a significant hurdle. iPSCs offer vast therapeutic and research potential, while more than 40 different monogenic disorders affect skeletal muscle, leading to progressive degeneration and atrophy—conditions for which effective treatments are still lacking ^26^. Skeletal muscle contains a tissue-specific population of muscle stem cells (MuSCs) that play a critical role in muscle regeneration, making them ideal candidates for cell replacement therapies ^27^. In the case of autologous applications within muscular dystrophy, the disease-causing genetic defect needs to be repaired before transplanting the cells back into the patient. Efficient and safe non-viral ex vivo delivery of genome editing tools is an essential first step toward achieving this goal.

In this study, we characterized and optimized a family of CPPs, known as hPep, that have previously been used for the delivery of nucleic acids ^28^. Here, we focused on adapting the hPep platform for efficient RNP-based delivery of Cas9 and its derivatives. During particle formation, enhanced delivery efficiency was achieved by incorporating silica as a core to the nanoparticle. This allowed for highly efficient delivery across cell types at low doses and delivery of more challenging cargos, such as prime editor and corresponding pegRNAs. hPep nanoparticles demonstrated virtually unaltered activity in serum conditions, irrespective of protein cargo. Furthermore, hPep enabled base editor delivery to primary cells such as MuSCs and iPSCs while preserving myogenic and pluripotency markers, respectively. The ability of hPep to efficiently deliver diverse cargos across various cell types represents a significant advancement towards a simple synthetic system. Importantly, this system can be easily prepared in minutes on a benchtop, stored as lyophilized powder, and applied using standard laboratory techniques.

## Materials and methods

### Cell-penetrating peptides

All peptides were procured from Pepscan Presto (Lelystad, The Netherlands) with a purity exceeding 90%. These peptides are terminated with C-terminal amidation and contain a free amine at the N-terminus. All peptides were supplied as lyophilized powder and solubilized in water before use. The peptide sequences are given in **Table 1**.

**Table 1.**
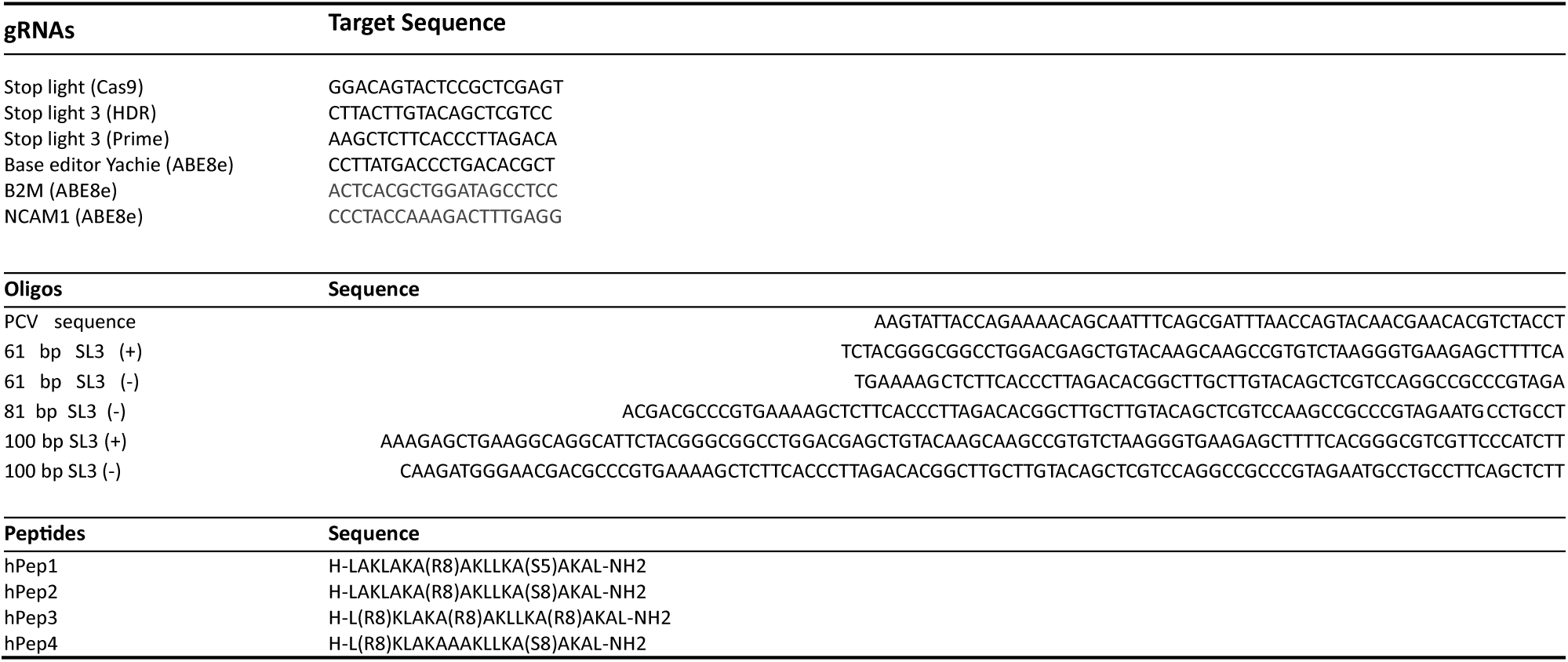
gRNAs, oligos, and peptide sequences.

### Reagents, chemicals, and media components

The Cas9 protein, CRISPR RNA (crRNA), trans-activating crRNA (trRNA), single-guide RNA (sgRNA), prime editing guide RNA (pegRNA), and oligonucleotides were procured from Integrated DNA Technologies (IDT), Coralville, IA, USA. Guide RNA sequences are available in **Table 1**. The crRNA, sgRNA, and trRNA contained IDT standard Alt-R™ modifications. The combination of crRNA (or xCRISPR RNA) with trRNA and the sgRNA will henceforth be called gRNA.

Hexametaphosphate, PEG8K, PVP10, PVP50, PVA18, PVA40, PVA-PEG (polyethylene glycol–polyvinyl alcohol, i.e., Kollicoat® IR), HEPES, 22 nm silica (LUDOX® TM-50 colloidal silica), Cell Proliferation Reagent WST-1 (version 18) Roche, and glucose was purchased from Merck, Sweden. The Pure PVA-PEG, an intermediate product in the Kollicoat® IR production, was given as a kind gift from BASF SE (67056 Ludwigshafen, Germany). Promega Firefly Luciferase Assay System (cat#16174), DAPI (cat# 62248), Lipofectamine™(RNAiMAX) (cat#13778075), Dulbecco’s modified Eagle’s medium (DMEM®) phenol red free (cat#31053028), DMEM Gluta-Max™ (cat#10564011), Trypsin-EDTA (0.05%), phenol red (cat#25300054) were purchased from Thermo Fisher Scientific (Waltham, MA, USA).

### Protein production

The His-NLS-Cre recombinase (herein called Cre), PCV-Cas9, Adenine base editor 8e (herein called ABE8e), and Prime editor 2 (herein called PE2) (Addgene plasmids: 62730, 123643, 161788, 132775 respectively), were expressed by Escherichia coli (BL21 (DE3) T1R pRARE2) upon induction at an optical density of 3 utilizing IPTG. Following induction, the proteins were purified using immobilized metal affinity chromatography using HisTrap HP (GE Healthcare), followed by gel filtration through HiLoad 16/60 Superdex 75 for Cre and Superdex 200 for the Cas9 fusion proteins. The fractions obtained from the gel filtration were assessed via SDS-PAGE gel electrophoresis before combining the main protein fractions. The purified proteins were stored in a solution comprising 20 mM HEPES, 300 mM NaCl, 10% glycerol, and 2 mM TCEP, with a pH value of 7.5. The proteins were produced and purified at the Protein Science Facility (PSF, Karolinska Institutet, Stockholm, Sweden). Note, the PE2 (Addgene #132775) was moved into the pNIC-CH2 plasmid, which modified the protein by adding a GS-linker and HIS-tag in the C-terminus. Experiments using ABE8e were performed using protein either from the PSF at KI or provided by Dr. Anja Schütz (Max Delbrück Center Protein Production and Characterization unit).

### Cell lines and cell culture

HeLa, T47D, B16F10, and MSC Traffic light cells were generated in-house using a lentiviral system with the Addgene plasmid #65726. The HEK293T, MDA-MB-231, and MCF-7 Stoplight cells, used to evaluate Cas9 RNP delivery, were generated as previously described ^29,30^. Nozomu Yachie kindly provided the ABE-GFP reporter HEK293T cell line (pLV-CS-121), while the N2A and B16F10 ABE8e reporter cells were generated similarly to the traffic-light cells using the Addgene plasmid #131126 ^31^Unless otherwise stated, Cells were cultured in complete media, DMEM Gluta-Max™ (10% FBS + 1% anti-anti), in a humified incubator at 37 ⁰C with 5% CO2. Cells were regularly split using trypsin-EDTA (0.05%). Karolinska Institutet iPSC Core facility, Stockholm, kindly provided human iPSC lines CTRL-5, CTRL-7, CTRL-9, and CTRL-10. The cells were cultured in mTeSR™ Plus (Stemcell technologies) and seeded as 20,000 cells/cm^2^ on 0.5*μ*g/cm^2^ iMatrix511 (Takara Biosciences) in a CO_2_ incubator at 37°C. 5*μ*M Y-27632 dihydrochloride (Tocris) was added during the first 24h of the split. Complete media replenishment continues daily with pre-warmed complete mTeSR^TM^ plus media until cells are ready for the next split. Cells were harvested as single cells with 1X TrypLE select (Gibco) during splits and counted using the Countess cell counter (Thermo Fisher Scientific).

Human MuSC isolation was performed as previously described ^32^. Immediately after the biopsy procedure, the muscle specimen was transferred into Solution A for transport (30 mM HEPES, 130 mM NaCl, 3 mM KCl, 10 mM D-glucose, and 3.2 μM Phenol red, pH 7.6). The fresh muscle specimen was manually dissected into fragments and then subjected to hypothermic treatment at 5°C for 4–7 days before downstream processing for MuSC isolation. After hypothermic treatment, fragments were further mechanically dissected, and small fragments were cultured in individual vessels in Skeletal Muscle Cell Growth Medium (SMCGM, Provitro) supplemented with 10% FCS in a humidified incubator with 5% CO2 at 37°C to allow outgrowth of oligoclonal MuSC colonies. The outgrowing colonies were expanded and characterized prior to cryopreservation. MuSC populations used in this study are listed in **table 2**. For experiments, primary MuSCs were cultured in basal SMCGM enriched with supplement mix (Provitro) in a humidified incubator with 5% CO_2_ at 37°C. For passaging, MuSC were washed with Dulbecco’s phosphate-buffered saline (DPBS) (Thermo Fisher Scientific) and detached with 0.25% Trypsin-EDTA (Provitro) or TrypLE Express (Gibco) at 37°C for 5 minutes. The medium was switched to Opti-MEM™ I Reduced Serum Media (Thermo Fisher Scientific) once the cells reached confluence to induce fusion into multinucleated myotubes.

**Table 2.** MuSC donors used in this study.

| Donor ID | Age at biopsy (y) | Gender (m/f) |
| --- | --- | --- |
| #1 | 39 | f |
| #2 | 25 | m |
| #3 | 50 | m |

### Protein–peptide nanoparticle formation and cell treatment

The protein was added to the buffer tested, followed by the addition of the CPP. This was rapidly followed by vortexing. The protein amount was kept constant at 10 ng/µl, with the CPP being varied according to the final tested molar ratio (MR) between the protein and the CPP. This was done in PCR tubes due to their low binding properties. The complex was allowed to form for 40 minutes before adding it to the cells. When silica was used, it was directly added into the buffer before adding any protein, at which point the procedure followed. Care was taken not to leave the diluted silica in a buffer not explicitly made for silica storage longer than necessary. For the Cas9 and Cas9 derivatives, the protein was complexed with gRNA at an MR of 1:1.1 protein to RNA. xcrRNA/trRNA was mainly used, but cr/trRNA and sgRNA were also used. No difference in efficiency was seen between the species, so the species were used interchangeably in this study. PegRNA was used for prime editing and treated as a sgRNA for complexation with prime editor 2.

Particles were added to cells, seeded 10K/well 1 day prior, in complete media (10% FBS with 1% anti-anti) in a 96-well plate unless otherwise stated. The molar ratio used between RNP and CPP was 1:125 unless otherwise noted. The particle-containing media was either left on the cells for the full duration of the experiment or exchanged for fresh media at 2 h (as stated in each figure). Exceptions to this are indicated in the relevant figure. Cells were analyzed between 4h and 3 days after the addition of the complexes based on the experiment in question (generally 3 days for Cas9 and prime editing treatments and 2 days for ABE8e). HEK293T was used for Cas9, ABE8e, and prime delivery evaluation unless otherwise stated.

The additives were tested by diluting 15 w/v% of additive (dissolved in H2O) in HBG to the desired w/v%. Further testing with PVA-PEG was done by diluting a 10% PVA-PEG (10 w/v% PVA-PEG, 6.3 w/v% sucrose dissolved in H2O under heating at 80⁰C until dissolved) solution with the desired base solution, often DMEM. The norm for this dilution is 1:1 PVA-PEG 10 w/v% with DMEM forming a 5 w/v% PVA-PEG (3.15 w/v% sucrose) solution.

The iPSCs were treated similarly to the cell lines described above. Briefly, ABE8e:hPep3 particles were added to iPSCs seeded a day prior as 10K/well in a 96-well flat bottom TC plate in 100ul. Before particle addition, cells have been given fresh pre-warmed mTESR Plus with 1X Anti/Anti (Gibco). Complete media was replaced 6 hours after treatment, followed by another media replacement 1 day after treatment. Cells were stained and analyzed by flow cytometry 3 days after treatment.

For NCAM1 targeting in HEK293T cells, 90K/well were seeded in 12-well plates 1 day before ABE8e RNP:hPep3 treatment. The media was changed at 4 h after treatment. Cells were harvested for analysis on day 2. For experiments in MuSC, 30K/well were seeded in 12-well plates 1 day before ABE8e RNP:hPep3 treatment. The media was changed after 6 h or overnight incubation (∼16 h), depending on the concentrations added. Cells were harvested for analysis on day 3 after treatment unless otherwise stated.

### RNAiMAX positive control RNP transfection

Transfection using RNAiMAX was conducted according to the protocol optimized by the Chesnut lab ^10^. These transfections were performed in complete media containing 10% FBS. Unless otherwise stated, editing efficiency was investigated on day 3 after treatment.

### Flow cytometry

Cells were analyzed 4 h to 3 days after treatment to investigate reporter expression. Cells were prepared for the flow cytometry by a PBS wash, followed by trypsin-EDTA (0.05%) digestion. They were stained with DAPI (Thermo Scientific, 25 ng/mL) before analysis by flow cytometry on a MACSQuant Analyzer 10 or 16 (Miltenyi Biotec, Bergisch Gladbach, Germany). Data from this experiment were analyzed using FlowJo 10.6.2 software (FlowJo, LLC).

Antibody staining was done by adding a titrated amount of labeled antibody to single cells, prepared as above, followed by incubation at +4⁰C and a wash with PBS to remove any unbound antibody. The live dead stains were performed using DAPI, except for the iPSC, where LIVE/DEAD™ Fixable Aqua Dead Cell stain (L34966, Invitrogen) was used as per manufacturer’s instructions after harvesting the cells with TrypLE select. The iPSCs were counterstained with Anti-SSEA4 (1:50, 561565, BD Pharmingen) antibody while staining with B2M (Beta-2 Microglobulin) (1:200, 316312 BioLegend) antibody for 60 min in the dark at +4⁰C to identify pluripotency state and B2M KO of the cells before acquiring on flow cytometer.

### Storage Testing

For vacuum concentration, 50 µL of the prepared complexes were transferred into 1.5 mL microcentrifuge tubes and dried under vacuum conditions using a Thermo Scientific Savant SC210A SpeedVac. The process was conducted without the application of additional heat. The dried samples were resuspended in 50 µL of H₂O before use.

For freeze-drying, 50 µL of the complexes were initially frozen at −80 °C and then transferred into a pre-chilled cooling block maintained at −80 °C. The samples were subjected to vacuum drying until complete desiccation. The lyophilized samples were rehydrated with 50 µL of H₂O before use.

Freeze-thaw cycles were performed by alternately freezing the samples at −80 °C and thawing them at 37°C.

### WST-1 assay

10 µl of the WST-1 assay reagent (Roche) was added to each well (96-well, 100 µl cell media) and was then incubated between 1-2h. Plates were measured before absorption values went above 2. NT, media only, and lysed cells were used as controls for background and normalization purposes.

### Gal9 imaging experiments and quantitation

Cells were seeded in MatTek glass-bottomed dishes (60,000 cells/35mm dish) in complete media (DMEM containing 10% FBS) a minimum of 16 h before treatment. Nuclei were uniformly stained with Hoechst 33342 (0.5 μg/ml) added to the culture medium 30 minutes before imaging experiments. Subsequently, the cell media was replaced with treatment-containing media at selected doses, and the Petri dish was promptly transferred to an inverted Nikon C2+ confocal microscope equipped with a humidified imaging chamber set at 5% CO_2_ and 37 °C.

Live-cell experiments were conducted using a 60× 1.4 Nikon APO objective and with C2-DUVB GaAsP detectors and variable emission bandpass, employing a 405 nm and 561 nm laser for relevant fluorophores. Images were processed and analyzed using Cell Profiler (v.4.2.6) image-analysis software. Hoechst 33342 facilitated nuclei detection, while diffuse mCherry-GAL9 fluorescence aided in cytoplasm identification. Punctate structures representing ruptured endolysosomes were identified through maximum-intensity projection fluorescence images.

Quantification of spot populations utilized the ’identify primary objects’ function within Cell Profiler, discerning punctate structures with intensities surpassing local background cellular intensity and ranging in size from 2 to 35-pixel units. Experimental replicates were averaged from varied individual acquisitions. The resulting data were exported and graphically represented using Origin 2022b.

### Genomic DNA (gDNA) extraction and PCR amplification

Genomic DNA (gDNA) was extracted using Agencourt AMPure XP beads (Beckman Coulter). Briefly, samples were lysed in a heating block at 56 °C for 10 minutes using AL Buffer (Qiagen) supplemented with 0.2 mg/mL Proteinase K (Qiagen). Following lysis, double the sample volume of prewarmed AMPure XP beads was added, and the mixture was mixed on a rotational wheel. The tubes were placed on a magnetic rack to separate the bead-bound gDNA from the supernatant. The beads were washed twice with 80% ethanol to remove contaminants, and the bound DNA was eluted with FG3 buffer (Qiagen).

The target region was amplified using Q5 High-Fidelity DNA Polymerase (New England Biolabs). The primer sequences used for amplification are listed in **Table 3**.

**Table 3.**
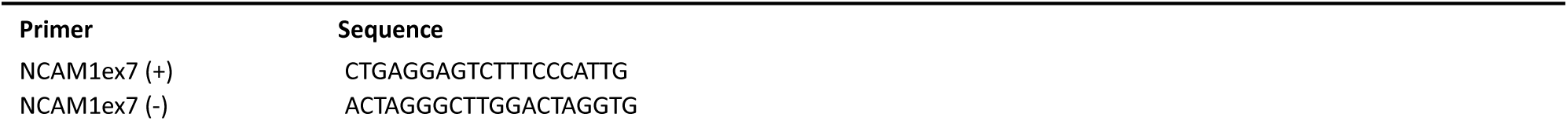
Primer sequence for NCAM1ex7 PCR amplification.

### Genome editing analysis by Sanger and Next-Generation Sequencing

PCR products were purified using the NucleoSpin Gel and PCR Clean-up Kit (Macherey-Nagel) according to the manufacturer’s protocol. Sanger sequencing was carried out by LGC Genomics (Berlin, Germany), and chromatogram analysis was performed using EditR software (v1.0.10) ^33^. For Next-generation amplicon sequencing, samples were analyzed at GENEWIZ (Amplicon EZ service) using an Illumina MiSeq platform and 250 bp paired-end reads. Results were analyzed using CRISPResso2 ^34^. The following parameters were applied: Sequencing design: Paired-end reads; Minimum homology for alignment to an amplicon: 60%; Center of the quantification window (relative to the 3’ end of the provided sgRNA): -10; Quantification window size (bp): 10; Minimum average read quality (phred33 scale): >30; Minimum single bp quality (phred33 scale): No filter; Replace bases with N that have a quality lower than (phred33 scale): No filter; Exclude bp from the left side of the amplicon sequence for the quantification of the mutations: 15 bp; Exclude bp from the right side of the amplicon sequence for the quantification of the mutations: 15 bp.

### Gel visualization of RNP-CPP binding (gRNA mobility assay, Hexametaphosphate)

Cas9 and cr/trRNA-Atto550 were complexed 1:1.1 MR, as mentioned above. The RNP was then complexed with CPP in different buffers for 40 min, and a total of 1 pmol RNP was loaded onto a 1% Agarose gel, which ran at 90 V, 200 A for 30 min. The sample was visualized using a Molecular Imager® VersaDoc™ MP 4000 system. Glycerol was added to a final concentration of 5% in samples formulated in any buffer not containing PEG or PVA-PEG to ensure complete loading into the agarose gels. The hexametaphosphate challenge was done by adding increasing amounts of hexametaphosphate for 5 minutes before loading it onto the gel, as described above.

### Encapsulation

Atto550-RNP was formulated and complexed with CPP, as mentioned earlier. One difference is the use of phenol-red-free DMEM during the experiment. After nanoparticle formation, the complexes were spun at 20k x g for 30 min. After this, the supernatant was collected, and the pellet was resuspended in the relevant buffer. Both fractions were measured either on the SpectraMax i3x (Molecular Devices, San Jose, CA, USA) or on the CLARIOstar (BMG LABTECH, Ortenberg, Germany) for atto550 content.

### Spin down testing

RNP-CPP particles were formulated as previously described and added to cells in a flat-bottom 96-well that was spun at 500 g or 2000 g for 30 min RT. The cells were grown in normal conditions for 3 days before analysis by flow cytometry.

### ZetaView

To determine particle size and concentration of the particles, a PMX-230 ZetaView TWIN instrument (Particle Metrix GmbH, Inning am Ammersee, Germany) was used with the corresponding ZetaView NTA software (8.05.16 SP3) for data analysis. Samples were diluted with HBG buffer before measurement. The following measurement settings were utilized: sensitivity 75, shutter speed 130, and frame rate at 30 frames per second.

### Field emission scanning electron microscope (FE-SEM) and energy-dispersive X-ray spectroscopy (EDS)

The structure and morphology of the freeze-dried samples were analyzed using a field emission scanning electron microscope (FE-SEM; Hitachi S-4800, Japan). The samples were mounted on conductive carbon tape using an accelerated 1-3 kV voltage and a current of 10 μA. The elemental composition of the materials was determined from 3 randomly selected areas for each sample, using the same field emission scanning electron microscope as the electron beam source using an accelerated voltage of 20 kV and a 10 μA current.

### Immunostaining and imaging

Human MuSC were plated on 8-well µ-Slides (ibidis, Germany). For myogenic and proliferation marker staining, 4,000 cells/well were seeded and fixed after 2 days. For differentiation, 10,000-12,000 cells were seeded per well as medium was switched to Opti-MEM™ I Reduced Serum Media (Thermo Fisher Scientific) once cells reached 70-80% confluence. Cells were then fixed after 3-5 days with a 4 % formaldehyde solution for 10 minutes at room temperature (RT). Cell permeabilization was performed using 0.2% Triton X-100, and blocking was done with 5% bovine serum albumin (BSA) in DPBS for 1 hour at RT. Primary antibodies were incubated overnight at 4°C in 1% BSA, as indicated in **table 4**. Alexa Fluor-conjugated secondary antibodies (Invitrogen) were incubated for 1 hour at RT (1:500 in DPBS). Nuclei were counterstained with Hoechst 33342 (Invitrogen). Images were acquired with a laser scanning confocal microscope LSM 900 (Carl Zeiss Microscopy), DMI 6000 fluorescence microscope, and Thunder Imager 3D microscope (Leica Microsystems) and processed with ZEN 3.4 Blue edition (Carl Zeiss Microscopy), Leica Application Suite X (Leica Microsystems), ImageJ (NIH) and Adobe Illustrator 2023. ≥200 nuclei were counted per sample to calculate percentage values for myogenic and proliferation markers. Fusion indices were calculated as the percentage of nuclei within myotubes (defined as ≥2 nuclei in one cell) versus the total number of nuclei captured at 20x magnification. A total of ≥200 nuclei were counted per condition.

**Table 4.**
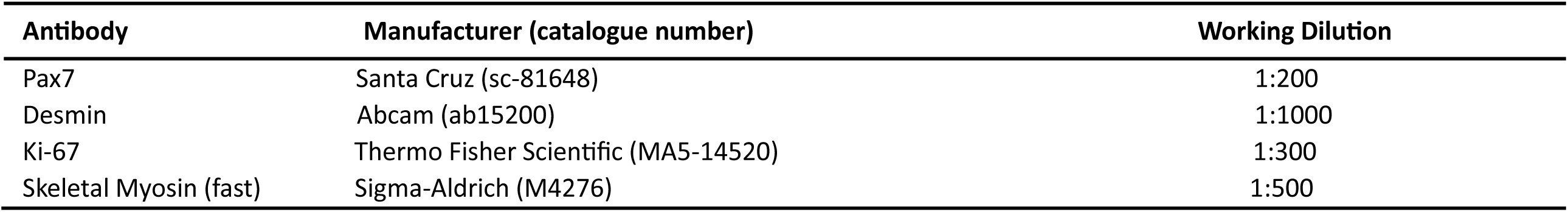
Primary antibodies that are used for immunostaining.

| Antibody | Manufacturer (catalogue number) | Working Dilution |
| --- | --- | --- |
| Pax7 | Santa Cruz (sc-81648) | 1:200 |
| Desmin | Abcam (ab15200) | 1:1000 |
| Ki-67 | Thermo Fisher Scientific (MA5-14520) | 1:300 |
| Skeletal Myosin (fast) | Sigma-Aldrich (M4276) | 1:500 |

### Statistical analysis

All statistical analyses were done using GraphPad Prism (9.2.0), with specific analysis methods described in the relevant figure legend. All error bars are SD, with each experiment being the means of three independent experiments unless otherwise stated.

## Results

### hPep-mediated delivery of Cas9-RNPs in reporter cells shows efficient delivery in full serum after formulation optimization

Cationic CPPs, initially designed for nucleic acid delivery, have been shown to form particles with Cas9 RNPs, facilitating intracellular delivery ^35^. To adapt the hPep family of CPPs for RNP delivery, four peptides were selected with identical amino acid sequences but varying alkenyl-alanine moieties, all with a total net charge of +6 (**Table 1**). The peptides were screened for their ability to deliver Cas9 RNP by combining the two components at different molar ratios (MR) followed by the addition to HEK293T Stop light (SL) reporter cells, which express GFP upon successful editing by Cas9 (**Fig. 1a**) ^36^. Among the candidates, hPep3 and hPep4 exhibited the highest delivery efficiency (**Fig. 1b**), whereas treatment with naked RNP resulted in editing below the detection limit of the assay, as shown by others ^30^.

**Figure 1.**
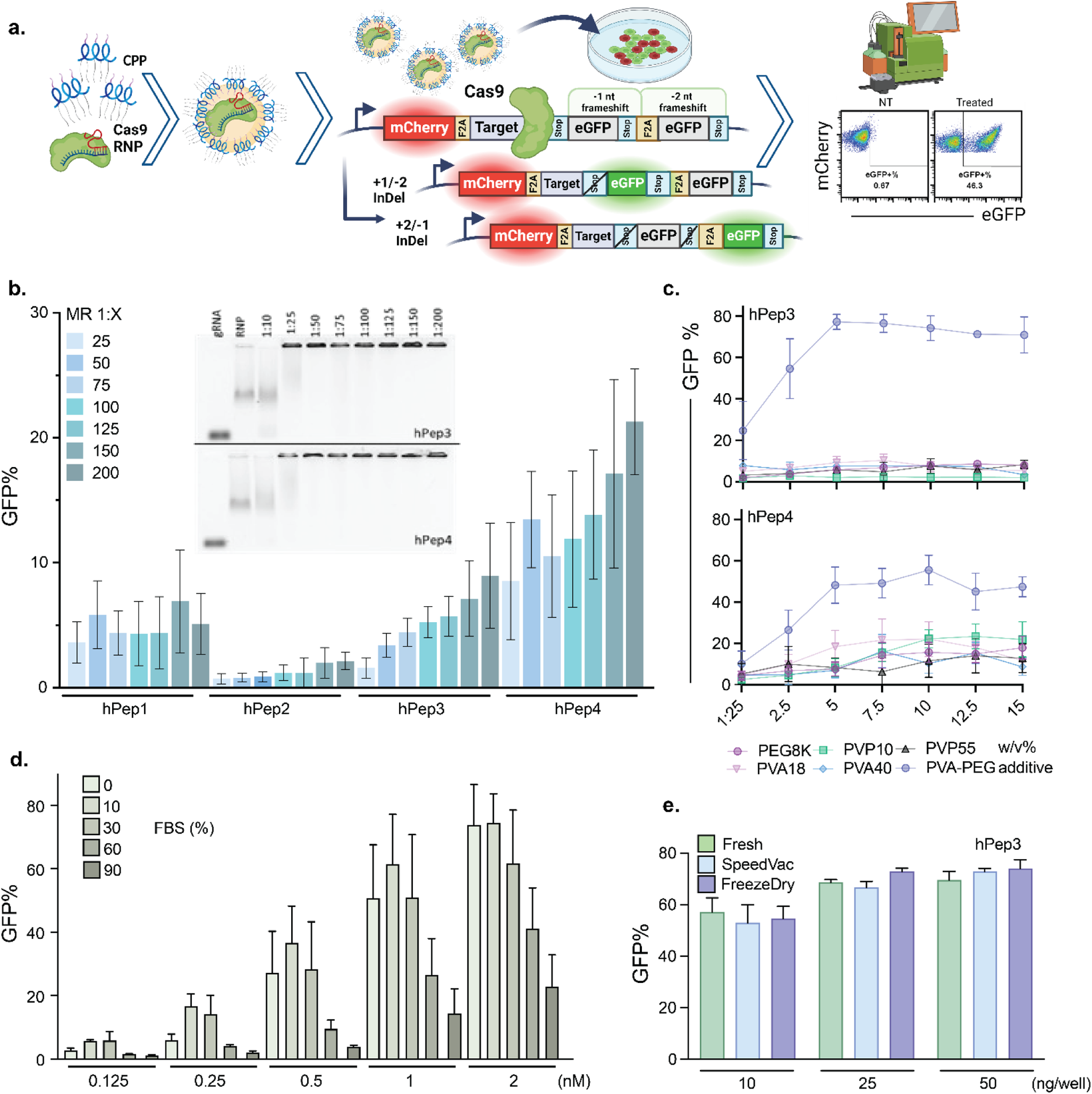
Experimental design and optimization of Cas9 RNP:hPep formulations in vitro. **a** Graphical illustration of the hPep particles, a schematic showing the Stoplight reporter system, and a general analysis method. **b** GFP% percentage after treatment with Cas9 RNP:hPep (200 ng Cas9 per 96-well, HEK293T SL cells) with increasing MR. n=4 independent experiments ± SD. Gel mobility assay to determine at what MR the RNP no longer migrates through the gel due to charge neutralization or particle size. **c** Screening of additives to enhance nanoparticle efficiency using hPep3 and hPep4 (25 ng of Cas9 per 96-well, HEK293T SL cells). The particles were formed in HBG diluted with the additive to a final concentration between 1.25-15 w/v%. Mean value of n=3 independent experiments ± SD. **d** Effect of serum on transfection efficiency of hPep3-CPP in PVA-PEG buffer (x-axis indicates conc. of Cas9 RNP added to HEK293T SL cells). One day after the transfection, serum was added to 10% in the 0% FBS condition. Mean value of n=3 independent experiments ± SD. **e** Testing of RNP:hPep3 compatibility with different storage methods (10/25/50 ng Cas9 per well. HEK293T SL cells). Mean value of n=3 independent experiments ± SD.

Next, various methods were used to characterize the formed nanoparticles, such as an electrophoretic mobility shift assay (EMSA), which confirmed nanoparticle formation at a low MR of 1:25 (**Fig. 1b**). However, optimal functional delivery was achieved at MRs 1:200 and above (**Fig. 1b**). A stability test using hexametaphosphate revealed that hPep4-formed particles were slightly more stable than those formed by hPep3, correlating with their higher transfection efficiency (**Supplementary Fig. 1a**). When particle formation was tested in different physiological buffers, HBG and Opti-MEM were the most effective, with HBG showing more consistent results between experiments (**Supplementary Fig. 1b**). Notably, spinfection of particles formed in HBG led to a ∼5-fold increase in editing efficiency (**Supplementary Fig. 1c**).

Nanoparticle tracking analysis (NTA) indicated a homogeneous population with a median size of 129 nm at 1:200 MR (**Supplementary Fig. 1d**). Particle size decreased from a median size of 192 nm at MR 1:25 to a stable range of 125-130 nm at MR 1:100 and above. Additionally, the number of particles larger than 200 nm substantially decreased with increasing MR.

The Cas9 RNP delivery efficiency achieved with the hPep family peptides was lower than with our previously published RNP-CPP delivery method ^35,37^. To address this, several additives were tested to enhance delivery efficiency (**Fig. 1c**). Among these, PVA-PEG showed the most significant improvement, reaching nearly 80% editing efficiency, surpassing previously published plasmid-based CRISPR/Cas9 transfection results using the same reporter system ^36^. It is important to note that the SL reporter system only activates GFP upon insertions or deletions of ±1 or ±2 base pairs, which inherently limits the maximum percentage of GFP-positive cells to below 100%.

To further optimize the delivery system, different buffers were combined with hPep3 and PVA-PEG during nanoparticle formation (**Supplementary Fig. 2a**). Using DMEM and Opti-MEM demonstrated the highest editing efficiencies, with DMEM being selected as the base buffer for subsequent experiments. Next, all hPep peptide variants were screened using the PVA-PEG/DMEM formulation, with as little as 10 ng (0.6 nM) Cas9 per well (**Supplementary Fig. 2b**). Interestingly, hPep1 and hPep2, which initially showed minimal to no transfection efficiency, now exhibited the highest editing efficiencies, reaching ≥75%, with hPep3 closely following.

Since hPep3 performed consistently well in both HBG and PVA-PEG buffer systems, it was selected as the lead candidate for further experiments. An EMSA assay revealed nanoparticles forming at lower MRs in the PVA-PEG buffer than in HBG, with minimal RNP smearing observed across all tested MRs (**Supplementary Fig. 2c**).

To assess editing efficiency in traditionally harder-to-transfect cell lines, MDA-MB-231 SL and MCF-7 SL cells were treated using RNP:hPep3 particles formulated at varying concentrations of PVA-PEG (**Supplementary Fig. 2**). As anticipated, editing efficiency was lower than that observed in HEK293T SL cells; however, it remained high.

To challenge the delivery system, increasing amounts of FBS were introduced to the culture media prior to transfection with Cas9 RNP:hPep3 particles (**Fig. 1d**, **Supplementary Fig. 2e**). Editing efficiency peaked at FBS concentrations ranging from 0% to 30% and gradually declined at higher FBS levels. However, editing remained detectable at 90% FBS. Additionally, the stability of the particles was evaluated under different storage conditions. The particles maintained their delivery efficiency after exposure to freeze-thaw cycles, vacuum concentration, or lyophilization (**Fig. 1e**, **Supplementary Fig. 2f & g**). These findings are consistent with our previous study, which demonstrated that RNP-CPP particles exhibit high resistance to challenging storage conditions ^35^. Thus, with the right additive, these peptides can efficiently deliver Cas9 RNP to different cell types in highly challenging conditions.

### hPep delivers protein cargos irrespective of charge and enables efficient co-delivery of HDR DNA templates

Traditionally, non-covalent CPP-mediated transfection relies on electrostatic interactions between cationic CPPs and anionic cargo. To investigate whether hPep particle formation is enhanced by increasing these electrostatic interactions, additional anionic charges were introduced to the cargo by coupling a single-stranded DNA (ssDNA) to the fusion protein PCV-Cas9, forming a covalent bond between these two components (**Supplementary Fig. 3a**) ^38^. However, adding the ssDNA did not improve editing efficiency (Fig. 2a). The oligo used was a random nucleotide sequence, except for the PCV binding site, designed to avoid genomic homology DNA binding, simply acting as an anionic charge carrier.

**Figure 2.**
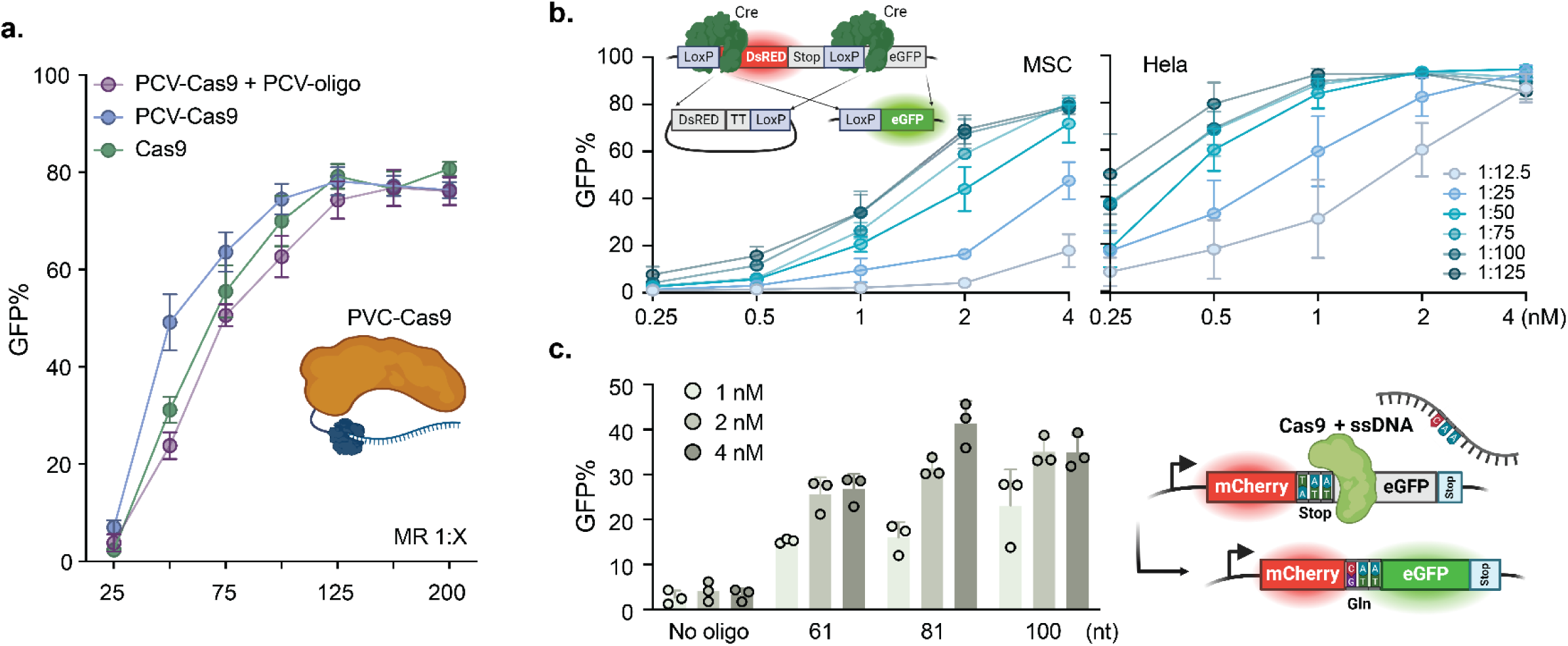
Cargo-independent hPep-mediated delivery and DNA-templated HDR editing. **a** Covalent attachment of a 60 nt oligo to PCV-Cas9 (53 nt after PCV binding). Particles were formed in DMEM/PVA-PEG buffer at increasing MR (15 ng protein per 96-well, HEK293T cells). Mean value of n=3 independent experiments ± SD. **b** Cre was complexed with hPep3 in DMEM/PVA-PEG buffer at increasing MR and added to MSC and HeLa cells harboring the TL reporter system in increasing concentration of Cre. Mean value of n=3 independent experiments ± SD. **c** HDR efficiency using a co-complexed ssDNA template. Particles were formed in DMEM/PVA-PEG (MR of 1:125) with increasing doses (1, 2, 4 nM) added to HEK293T harboring the SL3 system. Oligos were designed with homologous arms of equal length on each side of the expected DNA double-strand break site. Mean value of n=3 independent experiments ± SD.

To further investigate the relevance of cargo change, the cationic hPep was complexed with the cationic protein Cre recombinase. Interestingly, the peptides effectively delivered Cre to various cell types expressing a Cre-LoxP traffic light (TL) reporter system (**Fig. 2b**, **Supplementary Fig. 3b**). In some instances, nearly 100% recombination efficiency was achieved. Harder-to-transfect cell lines, such as T47D breast cancer cells and immortalized mesenchymal stem cells (MSCs), required higher MR and dosages to achieve higher editing levels, reaching up to 80%. These findings suggest that cargo charge does not impact delivery efficiency when using PVA-PEG, as strongly anionic and cationic cargo were both well tolerated in hPep3-mediated delivery.

Given the minimal effect of cargo charge on delivery efficiency, an attempt was made to co-deliver a homology-directed repair (HDR) ssDNA template to SL3 reporter cells. The ssDNA was designed to enable HDR-mediated conversion of a single thymine to a cytosine, thereby converting a stop codon to a glutamine codon (**Fig. 2c**) ^37^. Co-delivery of the HDR template resulted in ≥40% HDR rates using an 81 nt-long homologous ssDNA (**Fig. 2c**) ^37^. Notably, the choice of strand for the ssDNA template significantly influenced the editing outcome, with substantially higher HDR rates observed when the minus strand was used (**Supplementary Fig. 3c**).

### hPep-mediated delivery of gene editors does not confer cellular toxicity

Exploratory experiments pointed towards a quick mode of action, often coupled with adverse cell effects at high doses. A common solution for this is to shorten the cellular exposure time. Thus, the delivery kinetics of Cas9 RNPs using the hPep system were investigated in the SL reporter system, which requires intracellular uptake, endosomal escape, successful editing, and subsequent GFP transcription and translation for a positive signal to be detected. GFP expression was observed in treated cells as early as 4 hours post-treatment, indicating rapid uptake and efficient endosomal escape (**Fig. 3a**). Lipofectamine RNAiMAX exhibited significantly slower editing kinetics (**Fig. 3b**).

**Figure 3.**
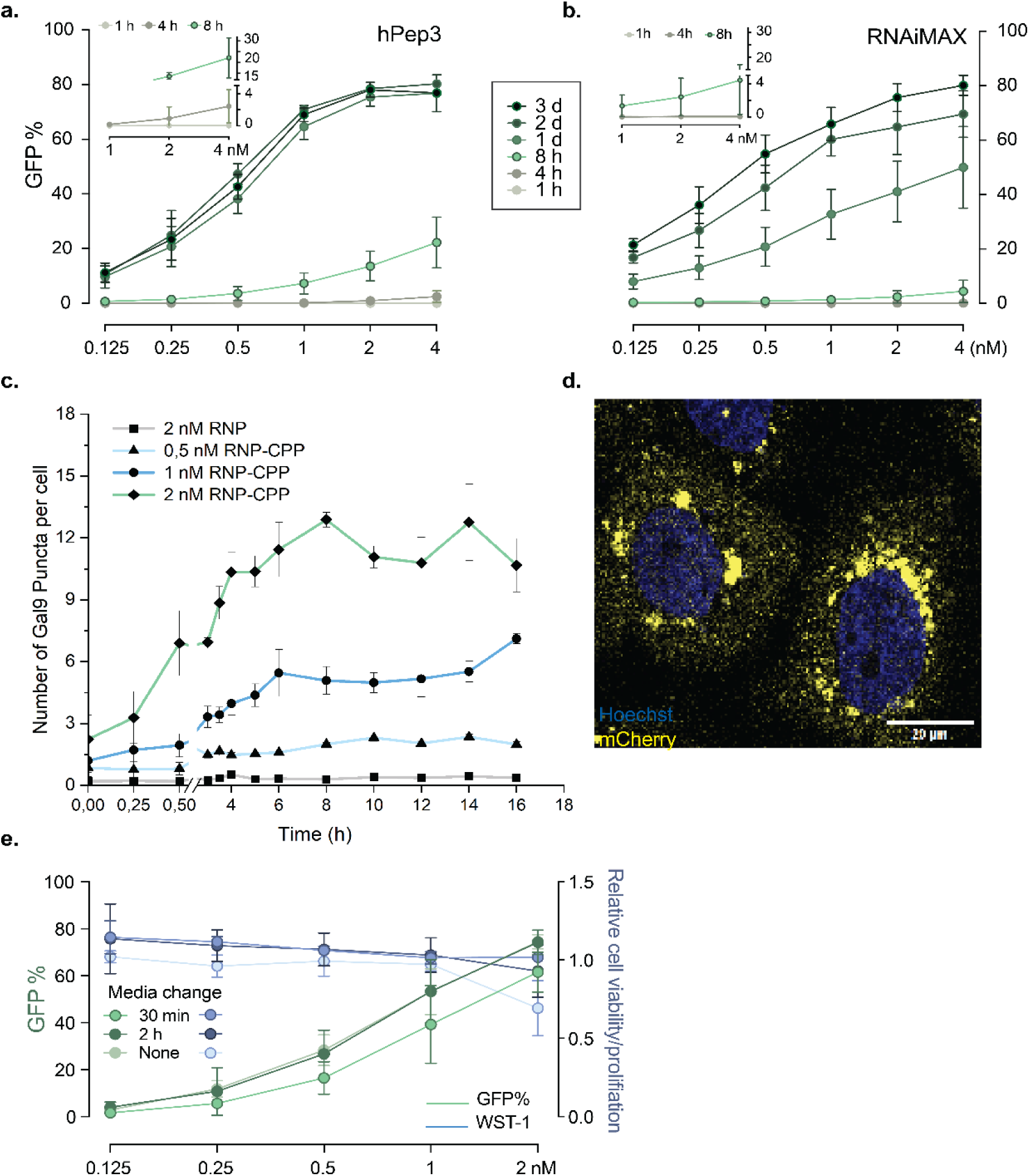
Rapid uptake and endosomal escape enable robust editing without associated cellular toxicity. **a &b** Editing rates after treatment with various doses of RNP:hPep3 (1:125 MR, HEK293T) or RNP:RNAiMAX. The treatment was performed simultaneously on 6 different plates, with each plate analyzed by flow cytometry at different time points. The RNP:hPep3 treatment resulted in detectable GFP at 4 h of treatment, reaching the maximum after 24 h. Editing with RNAiMAX was first detected at 8 h, increasing each day until day 3. The short-time points are magnified in the top left corner of a & b. Mean value of n=3 independent experiments ± SD. **c & d** A Huh7 mCherry-GAL9 reporter cell line was used to evaluate endosomal rupture. A rapid occurrence of rupture events after adding RNP:hPep3 formulated in DMEM/PVA-PEG was observed. n=3 independent experiments ± SD. The rupture events were observed to cluster in the perinuclear area. **e** A media change was implemented 30 min or 2h after adding the Cas9:hPep3. Editing rates were determined based on the activation of the GFP reporter (left y-axis in green), and cytotoxicity was measured using a WST-1 assay (right y-axis in blue) of identically treated cells. A 2-h incubation resulted in editing equivalent to a 3-day incubation, while cytotoxicity was almost eliminated at all tested doses. Mean value of n=3 independent experiments ± SD.

The swift delivery of Cas9-hPep3 was further corroborated in an endosomal rupture assay using a Huh7 mCherry-GAL9 (Galectin-9) reporter cell line ^39^. This reporter system relies on the recruitment of GAL9 to the site of an endosomal rupture, which can be quantified using the mCherry tag. Increasing concentrations of Cas9-hPep3 were added to the cells followed by microscopic analysis of mCherry-GAL9 puncta over 16 hours (**Supplementary Fig. 4a**). Puncta were detectable as early as 15 min after addition of nanoparticles, demonstrating rapid uptake and endosomal rupture with a plateau in puncta detection reached after 6 hours (**Fig. 3c**). Interestingly, the puncta were often concentrated in the perinuclear region, suggesting cargo release in proximity to the nucleus (**Fig. 3d**).

Quick and extensive endosomal rupture can lead to toxicity; hence, a WST-1 assay, measuring the metabolic activity of the cells, was conducted to assess the cytotoxicity of the particles. This revealed a dose-dependent reduction in metabolic activity following treatment with either hPep or RNAiMAX (**Fig. 3e**, **Supplementary Fig. 4b & c**). Given the rapid editing observed post-treatment, an evaluation was performed to determine whether a short exposure to the particles would suffice for efficient uptake and delivery while minimizing cellular toxicity. Indeed, a 15-minute incubation followed by a medium exchange yielded high editing levels while abolishing the previously observed toxicity (**Fig. 3e and Supplementary Fig. 4c**). A 2-hour incubation resulted in equivalent editing efficiency to conditions without medium exchange but with negligible toxicity and was hence chosen for subsequent experiments. In contrast, shorter incubation times drastically reduced editing rates when RNAiMAX particles were applied (**Supplementary Fig. 4d**). In summary, these data show that the hPep3-mediated transfection is exceptionally rapid and that adverse cellular effects can be negated by changing the media.

### Expanding hPep3– mediated delivery to base editors and prime editors

Recently developed Cas9 derivatives have demonstrated significant therapeutic potential, enabling the treatment of previously untreatable mutations. One such derivative is the base editor ABE8e ^40^. Therefore, we next wanted to assess if hPep3 is compatible with its intracellular delivery of ABEs. hPep3 and ABE8e RNP were complexed and introduced to HEK293T cells containing an integrated reporter cassette, designated HEK293T-ABE-GFP, where GFP expression is activated upon the conversion of a stop codon into a glutamine codon (**Fig. 4a**) ^31,40^. Increasing doses of ABE8e-hPep3 particles, formulated at varying MR, were administered to HEK293T-ABE-GFP cells, resulting in editing rates of approximately 90% even at the low dose of 1 nM RNP (**Fig. 4a**). These high editing rates were corroborated in B16F10 and N2A cells harboring the same reporter system, which demonstrated >90% editing efficiency (**Supplementary Fig. 5a)**.

**Figure 4.**
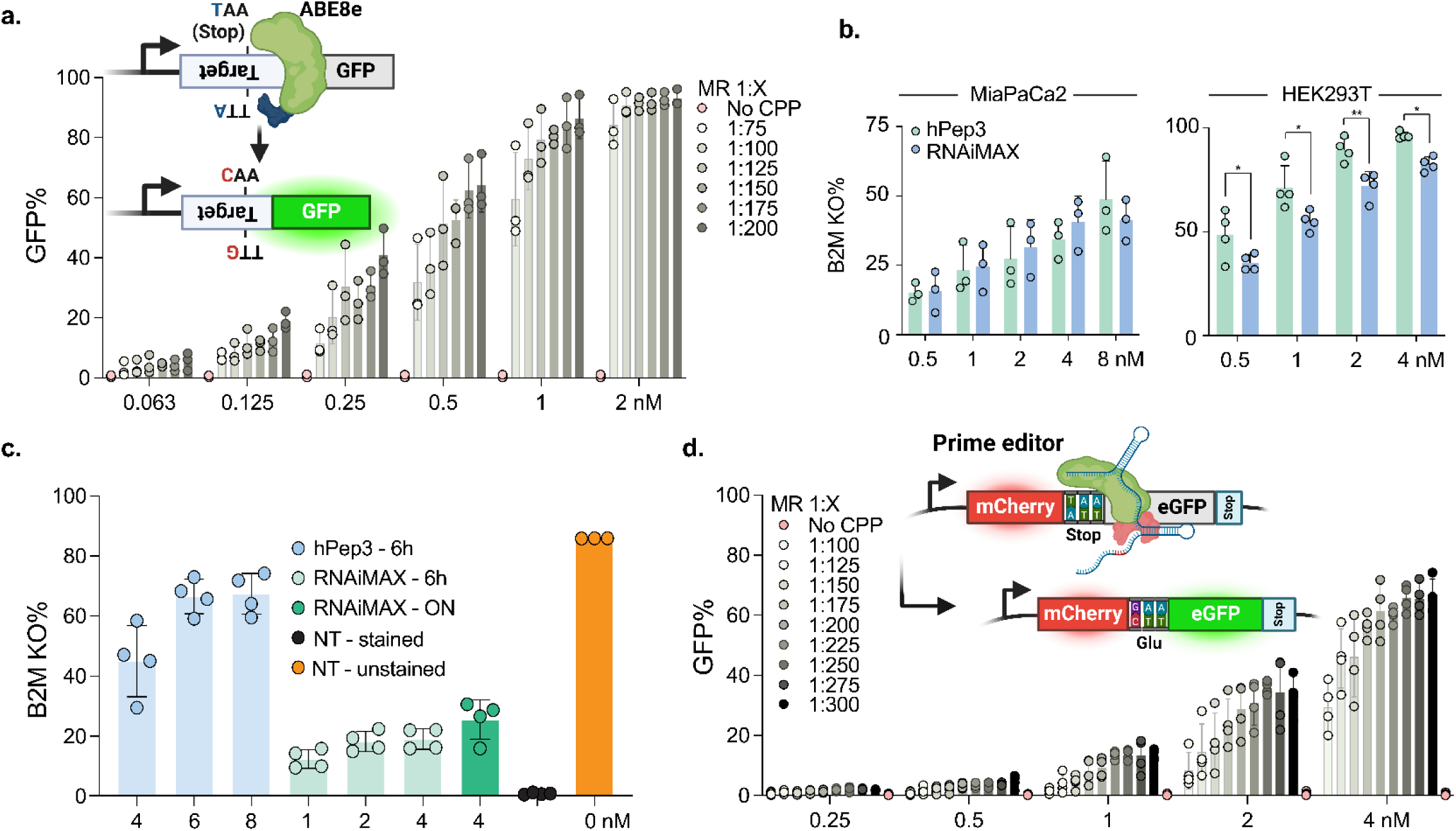
hPep3-mediated ABE8e and Prime editor delivery into different cell lines and iPSCs. **a** HEK293T-ABE-GFP cell was treated with an increasing dose and MR of ABE8e-hPep3 with a 2 h media change. Also shown is a schematic of the reporter construct and its mode of action. Mean value of n=3 independent experiments ± SD. **b & c** ABE8e mediated Knockout of B2M in MiaPaCa2, HEK293T WT cells, and iPSC by disrupting a splice donor site using hPep3 or RNAiMAX. The cells were stained with a B2M-APC antibody and gated for negative cells 3 days after treatment. The Y-axis displays % B2M negative cells. The NT cells were 100% positive for B2M (data not shown for MiaPaCa2 and HEK293T). The hPep3 and RNAiMAX groups were analyzed using 2way ANOVA and multiple comparisons, with p<0.05. Mean of n=3 (MiaPaCa2), n=4 (HEK293T) independent experiments ± SD. Human iPSCs from 4 different donors were treated with independent treatments performed in triplicates. Data is shown as mean value ± SD. Editing peaked at 86.4% for the highest edited repeat. A similar trend was observed for the other lines, higher than RNAiMAX at all tested concentrations. **d** Editing efficiencies of increasing doses and MR of PE2 given to HEK293T SL3 cells. Also shown is a schematic of the reporter construct and its mode of action. Editing reached 67%, with certain triplicates reaching 72% editing. The mean value of n=4 independent experiments ± SD.

Next, ABE8e was employed to induce mis-splicing of the pre-mRNA encoding the endogenous surface protein Beta-2 Microglobulin (B2M) by mutating a splice donor site causing a protein knockout (KO), as previously reported ^41^. Dose titration experiments were conducted using hPep3 and RNAiMAX to deliver the B2M-targeted ABE8e. Both modalities exhibited potent KO of B2M, with ABE8e-hPep3 achieving over 93% silencing, performing significantly better than RNAiMAX (**Fig. 4b**). The same delivery method was applied to iPSCs, resulting in B2M KO rates of up to 78% and 30% in hPep3-ABE8e and RNAiMAX treated cells, respectively (**Fig. 4c**). RNAiMAX was tested at lower doses due to associated cellular cytotoxicity, where the highest edited group, 4 nM ON (overnight), resulted in a 7-fold reduction in viable cells compared to the NT control. In contrast, the 6 nM ABE8e-hPep3 dose exhibited similar cell numbers and viability as the NT control three days post-treatment (**Supplementary Fig. 5b & c**). To further validate hPep as an iPSC delivery agent, cells were treated with a high dose of ABE8e:hPep3 (10 nM) for 6 h and stained for SSEA-4 (pluripotency marker) after 3 days. The cells maintained a high level of expression of the SSEA-4 marker (**Supplementary Fig. 5c**).

Another Cas9-derivate are prime editors, which hold tremendous therapeutic potential and enable precise gene manipulation such as the removal, insertion, or modification of tens of base pairs—capabilities beyond those of Cas9 and base editors ^42^. However, prime editors are among the largest Cas9 derivatives and lack efficient synthetic delivery systems. To address this, hPep3 was investigated as a delivery system for the PE2 variant. PE2 RNP complexes were formulated with hPep3 similarly to the ABE8e and delivered to SL3 HEK293T cells harboring a reporter cassette. This reporter system activates GFP expression upon successful prime editing by converting a stop codon into a glutamic acid codon, thereby inducing GFP expression (**Fig. 4d**). A dose-dependent increase in GFP-expressing cells in treated wells was observed, reaching up to 70% at the highest RNP dose and PE2:CPP molar ratio.

To the best of our knowledge, this is the highest editing achieved in iPSCs and the first instance of prime editor RNP delivery using a synthetic RNP delivery system.

### Colloidal silica enables protein absorption and is vital for high-efficiency RNP delivery

The fact that PVA-PEG addition in hPep formulations significantly improved editing efficiencies across all tested proteins, including the cationic Cre and large prime editors, prompted us to examine the role of the PVA-PEG additive. During the manufacturing process of PVA-PEG, colloidal anhydrous silica particles are added to aid in spray-drying of the PVA-PEG. Silica is known for its ability to adsorb proteins, peptides, and nucleic acids and has been used in various transfection methods, although none involving CPPs ^43–48^. Further investigation was warranted, given the presence of silica in the PVA-PEG and its potential role in enhancing the potency of hPep-formulated nanoparticles.

A gel electrophoresis showed that the Cas9 protein did not migrate into the gel when PVA-PEG was present (**Fig. 5a**). At the same time, a hexametaphosphate challenge indicated a weak interaction between the RNP and silica (**Supplementary Fig. 5d**). Silica was next isolated from the PVA-PEG solution by centrifugation and imaged using scanning electron microscopy, revealing particles in the sub-50 nm range (**Fig. 5b**). When silica-depleted PVA-PEG supernatant was used as a formulation buffer, a marked decrease in transfection efficiency was observed compared to non-depleted PVA-PEG (**Fig. 5c**). Energy-dispersive x-ray spectroscopy confirmed the presence of silica in the pellet and none in the supernatant (**Supplementary Fig. 5e**).

**Figure 5.**
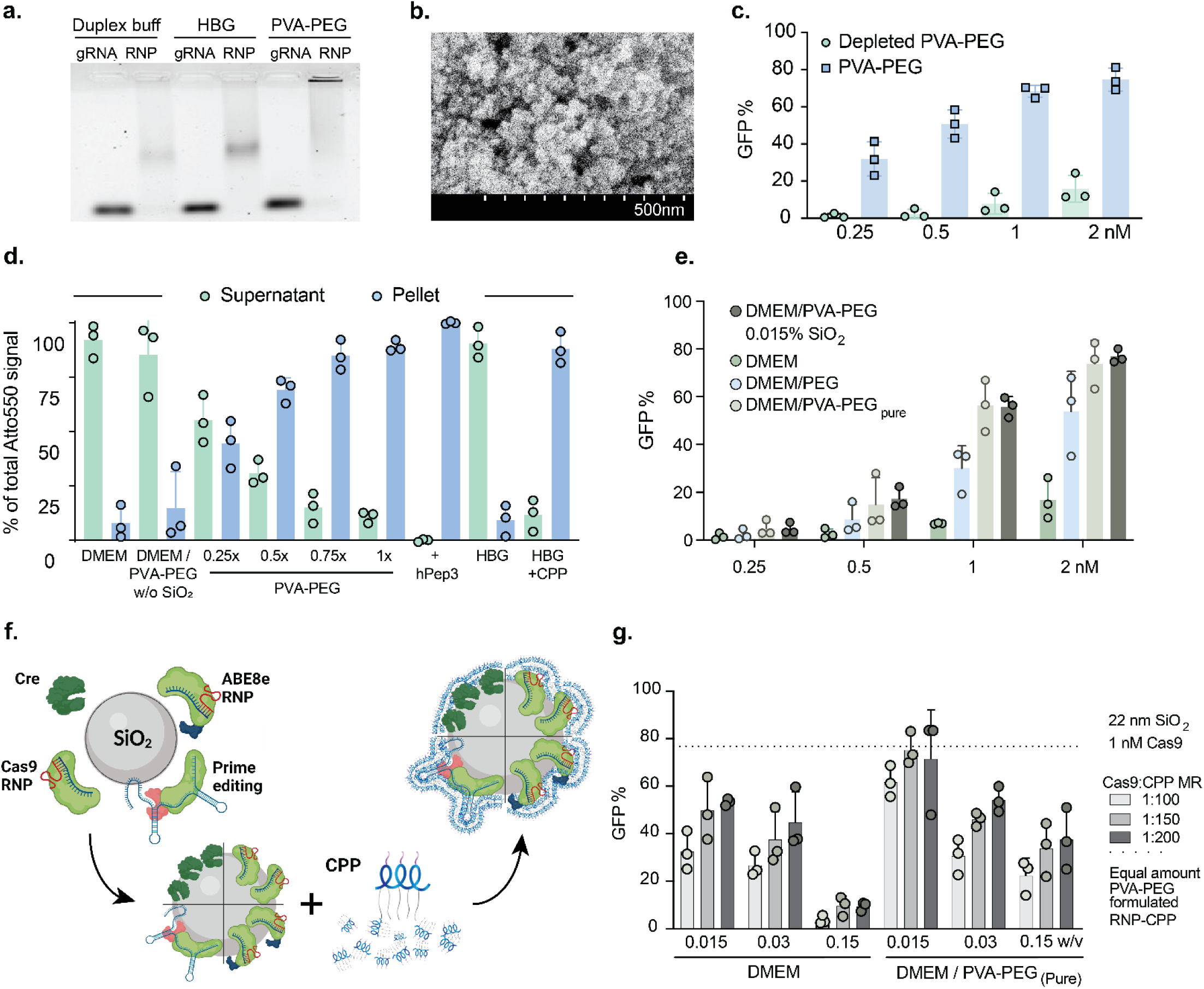
Exploration of the RNP-silica interaction and its effect on hPep-mediated delivery efficiencies. **a** Gel electrophoresis of Atto550 tagged gRNA or Cas9 RNP in different buffers. The RNP was unable to migrate through the gel when incubated with PVA-PEG. **b** FE-SEM of isolated silica particles from the PVA-EPG solution. Individual particle size appears to range from 20-30 nm. Each tick in the scale bar equals 50 nm. (**c**) Effect of removal of silica on transfection efficiency of HEK293T SL cells treated with hPep3-RNP. Mean value of n=3 independent experiments ± SD. **d** Atto550 tagged RNP was formed under different conditions fractioned by centrifugation, with % of Atto550 signal measured in each fraction. Mean value of n=3 independent experiments ± SD. **e** Silica isolated from the PVA-PEG buffer by centrifugation was added to different buffers, with the addition to DMEM/PVA-PEG_Pure_ recapturing the efficiency of RNP-CPP formulated in normal DMEM/PVA-PEG when added to HEK293T SL cells. Note: After dilution into the working buffer, the PVA-PEG contains 0.015% silica. Mean value of n=3 independent experiments ± SD. **f** Envisioned particle formation and structure. Results indicated that the protein adsorbed onto the silica surface, with the peptide interacting with both species. **g**) Commercial 22 nm Silica beads were added to DMEM and DMEM/PVA-PEG_Pure_ in increasing amounts compared to the PVA-PEG-derived concentration. Adding 1x, 0.015% w/v%, the concentration compared to PVA-PEG into PVA-PEG_Pure_ resulted in a similar efficiency as normal PVA-PEG when transfecting HEK293T SL cells. Mean value of n=3 independent experiments ± SD.

To further explore this effect, Atto550-tagged gRNA was complexed with Cas9 RNP and added to buffers with or without PVA-PEG or hPep3. After centrifugation, Atto550-RNP localized in the pellet fraction in a silica concentration-dependent manner (**Fig. 5d**). When hPep3 was added, 100% of the Atto550 signal was localized in the pellet, indicating complete encapsulation of the Cas9 RNP.

Next, the centrifugation-isolated silica was tested by adding it to DMEM, DMEM/PEG8K, and DMEM/PVA-PEGPure (PVA-PEG without silica) in increasing amounts, along with a fixed amount of RNP:hPep3. Optimal delivery was observed at silica concentrations matching those found in the PVA-PEG buffer (**Fig. 5e & Supplementary Fig. 5f**), with PVA-PEG contributing to the effect but to a lesser extent than silica. The envisioned mechanism involves protein adherence to silica, with the CPP binding to silica and RNPs (**Fig. 5f**).

Commercial 22 nm silica beads were subsequently used with Cas9 RNP in DMEM or DMEM/PVA-PEG_Pure_ to corroborate the silica effect found in PVA-PEG. Maximum editing was achieved when silica concentrations matched those of the PVA-PEG buffer, with results comparable to those of the PVA-PEG containing silica (**Fig. 5g**).

In conclusion, adding silica is vital for high-efficiency delivery using the hPep system in this configuration. Furthermore, the combination of silica and hPep3 resulted in a 100% encapsulation rate of Cas9 RNP. Lastly, while the silica used in this work was added to the PVA-PEG by the manufacturer, thus potentially reducing reproducibility, our work also shows that it can be replaced by colloidal 22 nm silica.

### Pep3 enables highly efficient and well-tolerated ABE8-mediated base editing of primary human muscle stem cells

In light of the encouraging base editing results achieved in reporter cells and iPSCs, hPep3-mediated base editing was next assessed in therapeutically relevant primary human MuSC. MuSC holds great potential in cell replacement therapies for muscle wasting disorders but has been traditionally considered difficult to transfect. The *NCAM1* gene encodes for neural cell adhesion molecule 1, a membrane protein ubiquitously expressed in human MuSC ^49^. We previously showed that the knock-out of *NCAM1* has no negative impact on MuSC fitness ex vivo. Therefore, this gene serves as a universal endogenous reporter locus to assess the efficiency of gene editing interventions in MuSC from any donor . We hence targeted a splice donor site in *NCAM1* exon 7 using ABE8e (**Fig. 6a**) ^50^. Increasing doses of *NCAM1*-targeting ABE8e:hPep3 nanoparticles were applied to MuSCs derived from three different donors with a 6-hour incubation. This resulted in ∼90% A>G conversion at the highest doses 3 days after the treatment, with minimal indel formation (**Fig. 6b**). Extending the incubation time to 16 hours further improved editing efficiency, particularly at lower doses, (**Fig. 6c**). The treatment caused a moderate reduction in cell growth during the first three days post-treatment, independent of the editing status. However, the proliferation rate normalized after 3-5 days, remaining comparable to untreated cells throughout the remainder of the observation period (**Fig. 6d and Supplementary Fig. 6a**). No significant differences were detected in the expression of key markers such as PAX7 (MuSC marker), Desmin (pan-myogenic marker), or Ki-67 (proliferation marker) between treated and untreated MuSCs at any ABE8:hPep3 dose (**Fig. 6e & g**, **Supplementary Fig. 6b**).

**Figure 6.**
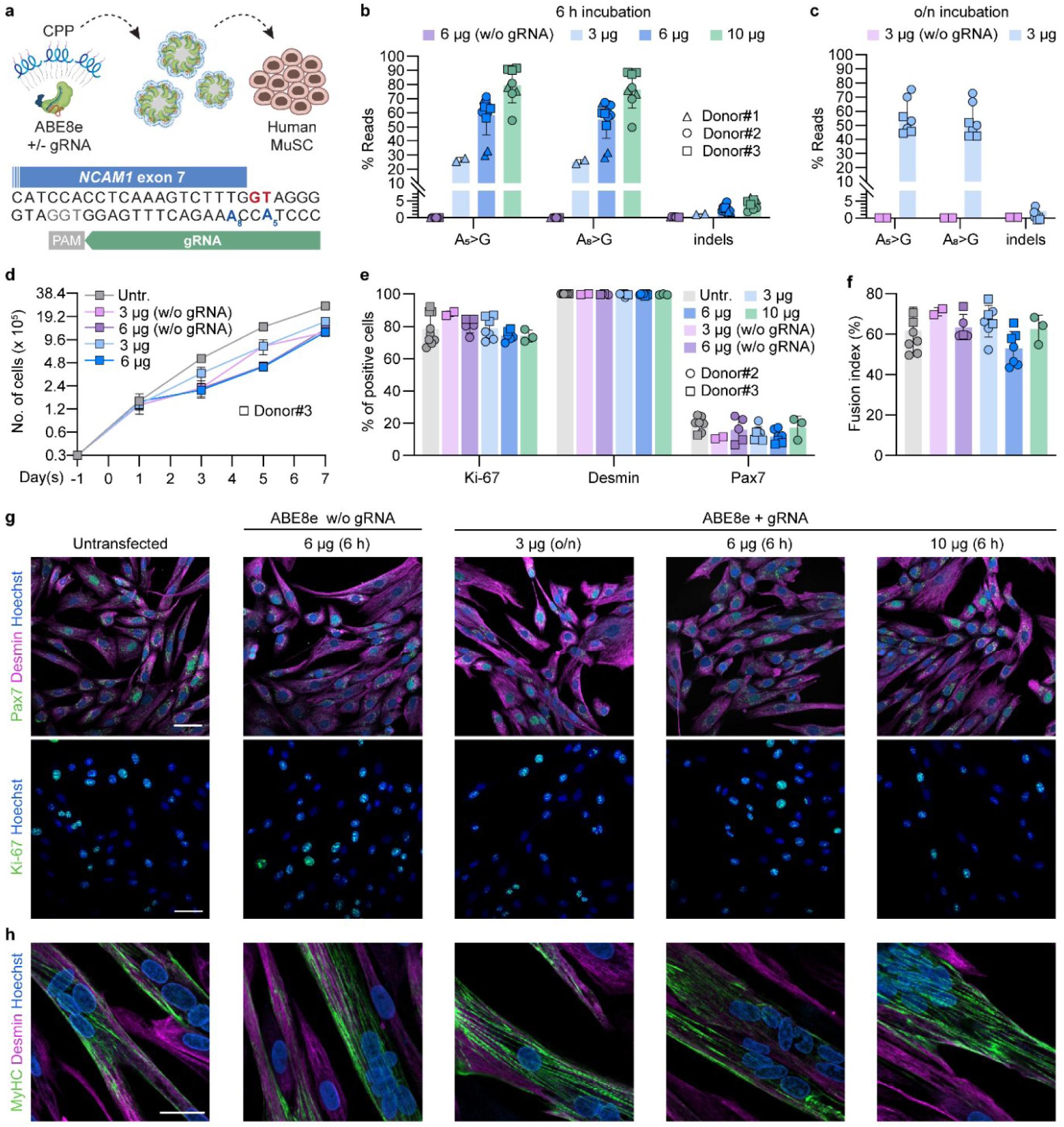
Editing efficiency, marker expression and morphology of human MuSC. **a** Schematic of the experimental set-up. gRNA targeting *NCAM1* exon 7 splice donor site (GT, red). Adenines located at protospacer positions 5 (A_5_) and 8 (A_8_) are highlighted in blue. The PAM sequence is shown in grey. Cells were grown in 12-well plates, harvested 3 days after treatment followed by processing for gRNA analysis. **b** Percentage of NGS reads with A>G conversion at A_5_ and A_8_ for MuSC from three donors after treatment with hPep3-ABE8e. **c** Percentage of NGS reads with A>G conversion after treatment with hPep3-ABE8e. **d** Growth analysis of human MuSC over 7 days after treatment with hPep3-ABE8e targeting *NCAM1* exon 7 (*n* = 2-3 technical repeats, mean ± SD). Cells were seeded at day -1 and treated at day 0. **e** Quantification of myogenic (Pax7, Desmin) and proliferation markers (Ki-67) at day 5 after treatment with hPep3-ABE8e targeting *NCAM1* exon 7 (*n* = 2-6 technical repeats per donor, mean ± SD, ≥ 200 cells/nuclei counted per condition. **f** Fusion indices calculated as a percentage of nuclei within multinucleated MyHC-expressing myotubes (*n* = 2-5 technical repeats per donor, mean ± SD, ≥ 200 nuclei were analyzed per condition). **g** Confocal microscopy images of MuSC from donor #2 immunostained for myogenic and proliferation markers after treatment with hPep3-ABE8e targeting *NCAM1* exon 7. Scale bars: 50 µm. **h** Confocal microscopy images of MuSC from donor #2 induced to fuse into multinucleated myotubes after treatment with hPep3-ABE8e. Scale bar: 50 µm. Untr.: Untransfected.

Treated MuSCs were next induced to fuse into multinucleated myotubes to assess their myogenic potential, mimicking in vivo muscle fiber differentiation. The treated cells retained their ability to form multinucleated myotubes and express the differentiation marker Myosin Heavy Chain (MyHC), showing no adverse effects of the ABE8e treatment (**Fig. 6f & h**, **Supplementary Fig. 6c).** Lastly, moderate editing efficiencies at lower ABE8e doses were significantly enhanced after a second round of treatment (**Supplementary Fig. 7**).

These findings demonstrate high editing rates and negligible side effects in these hard-to-transfect cells, suggesting the potential for clinical applications in treating degenerative muscle diseases. This highlights the broad therapeutic implications of the novel CPP formulations developed here.

## Discussion

Gene editing as a cure for genetic diseases relies on the efficient delivery of relevant effector molecules, such as Cas9-RNPs, which depends on the level of cargo protection, internalization, and endosomal release. This study optimized the hPep family of cell-penetrating peptides to deliver Cas9 RNP and its derivatives. A key finding in this study is that these peptides can successfully deliver a diverse range of challenging protein cargos, including Cre recombinase and prime editors. Despite its overall cationic charge, Cre recombinase, typically deemed unsuitable for delivery via cationic CPPs, was successfully delivered at editing rates approaching 100%. This underscores the cargo-agnostic potential of this delivery system. Prime editors pose a significant challenge due to their considerable size and the highly exposed pegRNA, yet successful, efficient delivery using hPep3 was achieved.

The high editing rates observed in this study can be partially attributed to the introduction of PVA-PEG in the formulation. This additive significantly enhanced editing efficiency by approximately two orders of magnitude compared to the HBG-formulated RNP. During the PVA-PEG production process, silica particles are added to aid the spray-drying process. Silica is known for its ability to adsorb various molecules, including Cas9 RNP, as demonstrated in **Fig. 5a**. This led us to the hypothesis that both the protein and peptide components adhere to the silica surface, with the peptides further interacting with the anionic gRNA (**Fig. 5f**). Our hypothesis is reinforced by the successful delivery of Cre, indicating that factors beyond electrostatic interactions between the cargo and the CPP are involved, namely that of silica in the PVA-PEG formulation. The use of silica as a core for protein and CPP adsorption is particularly significant, as it alters the dynamics and behavior of the resulting nanoparticles. While the application of silica in conjunction with Cas9 RNP is not entirely novel, its combination with CPPs has not been explored previously ^44–48,51,52^. Incorporating silica to facilitate protein and peptide adsorption could enable previously incompatible cargo and peptide combinations since CPP-mediated non-covalent protein delivery typically relies on the previously mentioned electrostatic interaction between cargo and CPP.

Prime editing, alongside similar techniques like click editing, is heralded as the future of gene editing due to its capacity to make substantial modifications without inducing double-stranded breaks ^42^. However, one of the significant challenges associated with prime editing is its substantial size, ≈242 kDa, which is approximately 80 kDa larger than SpCas9, and its highly exposed pegRNA. Consequently, delivering large protein-RNA complexes, such as prime editors in RNP form, poses substantial difficulties, particularly without employing physical methods such as electroporation ^53^. Currently, only two examples of nanoparticle-based delivery of prime editors exist. The first, conducted by the Mikkelsen lab, utilized lentivirus-derived nanoparticles to achieve around 6% editing in HEK293T cells using PEmax, an optimized variant of PE2 utilized in this study ^54^. The second example, published recently by the Lui lab, reported approximately 60% editing using engineered virus-like particles (VLPs) after significantly engineering the prime editor and pegRNA ^55^. Excitingly, hPep3 demonstrated remarkable efficacy, achieving PE2-mediated editing rates approaching 70%. This performance significantly outstrips the approximately 20% editing efficiency achieved through nucleofection of PE2 mRNA in the same cell line (**Fig. 4d**). However, it is important to note that different pegRNA targets and specific DNA edits can exhibit varying propensities for successful editing ^56^.

Additionally, recently published click editors, which operate in a manner similar to prime editing, utilize ssDNA as a template that interacts with a DNA polymerase ^57^. While it remains premature to predict which method will dominate the gene-editing landscape in the future, it is apparent that cell-derived nanoparticles, such as VLPs, are unlikely to facilitate the efficient delivery of click editors. This is primarily due to the absence of ssDNA in the cytoplasm, rendering it unavailable for incorporation into gene-editing vectors. Therefore, delivering click editors will likely require a fully synthetic approach akin to the methodology described in this article.

While the prime editing presented herein demonstrates significant potential, it is not the only RNP of interest explored in this study. Base editors, such as ABEs and CBEs (cytosine base editors), can correct specific nucleotide substitutions and knock out various genes by disrupting their splice sites. These two widely used base editors can potentially correct approximately 25% of human pathogenic single nucleotide polymorphisms (SNPs), with the ongoing development of new base editors likely to expand this range further ^58^. Thus, it is exciting that ABE8e is so efficiently delivered, with nearly 100% editing efficiency observed both in a reporter target and B2M targeted KO experiments (**Fig. 4a & b**, **Supplementary Fig. 5a**). The KO of B2M is relevant for allogenic non-HLA matched donations of cells to patients, such as in allogenic CAR-T treatments, where the disruption of B2M is one of the proposed initial steps for developing off-the-shelf allogenic CAR-T products ^59^. This study also shows the capacity of hPep to deliver Cas9 RNP along with DNA templates, achieving efficient HDR editing in cells with editing levels exceeding 40% without the addition of HDR-enhancing agents (**Fig. 2c**).

The editing of iPSCs is of great interest because of their ability to differentiate into any cell type and thus presents tremendous applications to the field for both research and therapeutic applications. However, current transfection methods lack efficient delivery and negatively impact cellular viability ^60^. Furthermore, no efficient synthetic methods exist for RNP gene editor delivery into iPSCs. In contrast, hPep3-mediated delivery achieves high editing levels (77%) at 6 nM ABE8e while maintaining cellular viability and pluripotency phenotype (**Fig. 4c**, **Supplementary Fig. 5b & c**). This is not the first example of nanoparticle RNP delivery to iPSCs, but to the best of our knowledge, it is the most efficient nanoparticle RNP delivery and the first example of synthetic nanoparticle base editor delivery to iPSCs ^61^.

Finally, the hPep technology demonstrates significant promise for delivering gene editors to primary human MuSCs, proving compatible with downstream applications that require high editing efficiencies while minimizing cytotoxicity. Notably, up to 90% editing was achieved while not adversely affecting the myogenic properties of the cells. The MuSC proliferation rate returned to normal within approximately three days, indicating that MuSCs are well-suited for therapeutic interventions using the hPep system. Comparable high editing efficiencies in human MuSCs have also been reported with mRNA-mediated delivery of ABE and Cas9 via nucleofection ^62,63^. However, unlike nucleofection, CPP-mediated delivery is not constrained by cell number requirements, facilitating earlier application and reducing the need for extensive cell passaging and expansion before gene repair. These findings offer promise for developing future MuSC-based gene editing therapies for currently untreatable diseases. A promising direction for such future research would involve repairing mutations in patient-derived MuSCs through hPep3-mediated delivery of various CRISPR enzymes, including prime editors. Additionally, our findings pave the way for novel therapeutic applications in the context of in vivo genome editing of skeletal muscle, an area where progress with non-viral delivery methods for CRISPR tools has been limited ^62^.

Together, these findings demonstrate significant potential for the efficient delivery of various protein cargos, which can be applied to a wide range of cell lines and primary cells. The reagents used are low-cost to produce and straightforward to handle, requiring no specialized equipment. Further research is essential to translate the combination of silica, hPep peptides, and gene editors into ex vivo and in vivo therapeutic applications.

## Author contributions

O.G., S.K., M.G., O.S, R.C., B.B, S.R, Y.E., X.L and J.R. performed experiments. O.G. carried out in vitro cell work, excluding the primary muscle stem cell work which S.K and H.E. carried out (treatment, DNA sequencing, immunostaining, and imaging, etc.) and the iPSC work that S.B carried out. S.R. carried out work with PCV-Cas9. M.G. carried out Gal9 assays. O.S. carried out zetaview. B.B. carried out electron microscopy and energy-dispersive X-ray spectroscopy. O.G. carried out encapsulation and gel retardation experiments. X.L carried out stable cell line generation. Study design and critical discussions were carried out among O.G., S.K., C.I.E.S., R.C., O.D.J., X.L, E.A, Si.S., H.E., J.Z.N., and S.E.A, J.Z.N, E.A, and Si.S. provided funding. S.E.A., J.Z.N, and H.E. provided project directions. The manuscript was written by O.G., J.Z.N, H.E., and S.E.A., with input from all authors.

## Acknowledgments

Si.S. is an inventor on a technology for primary human muscle stem cell isolation and manufacturing (IP: (DE10 2014 216872), 2015 PCT (WO 2016/030371), granted in EU and US). Si.S. and H.E. are co-inventors on a pending patent application on gene editing of human muscle stem cells (European Patent 666 Office 21 160 696.7). Si.S. is co-founder of MyoPax GmbH and MyoPax Denmark ApS.

---

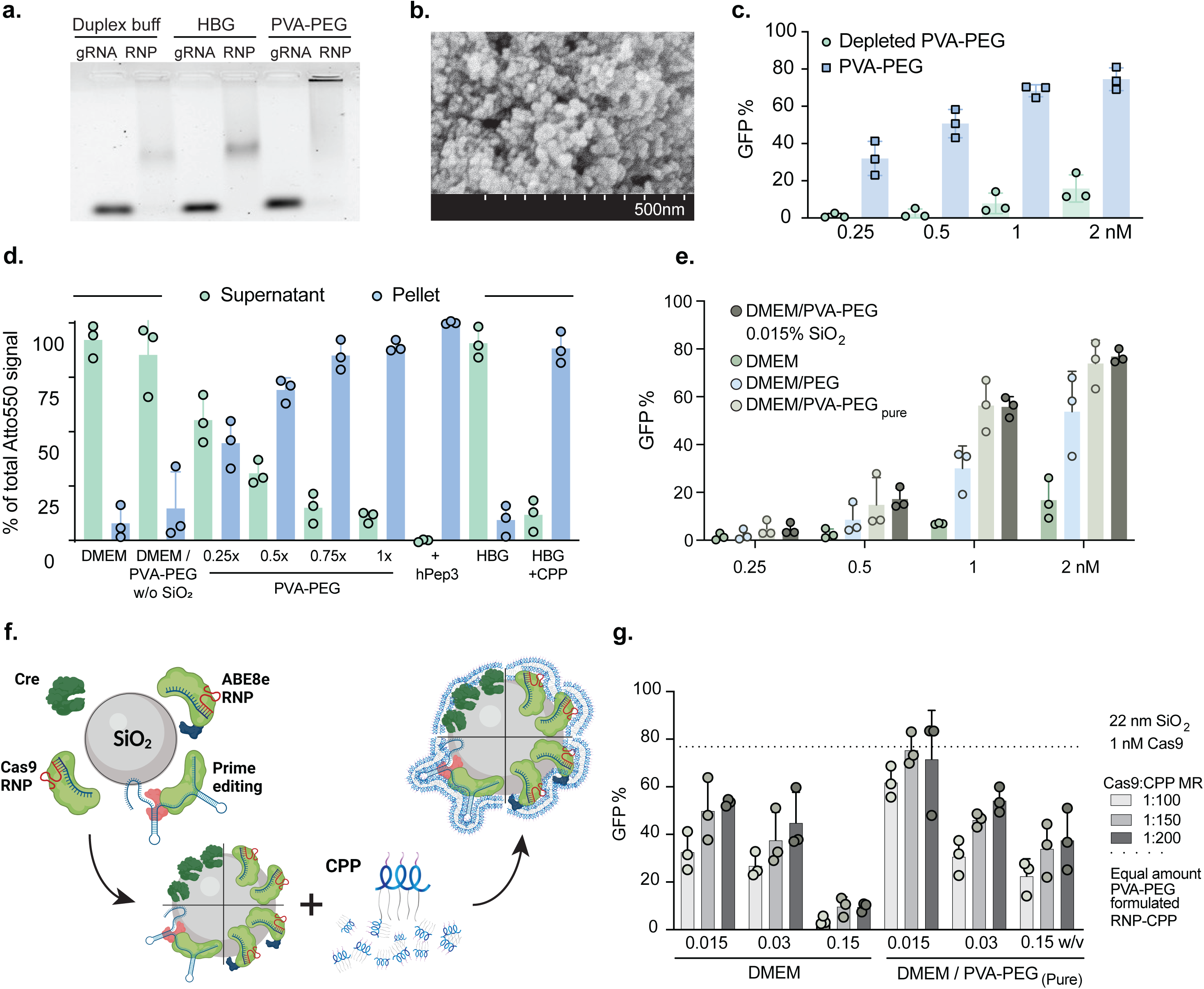

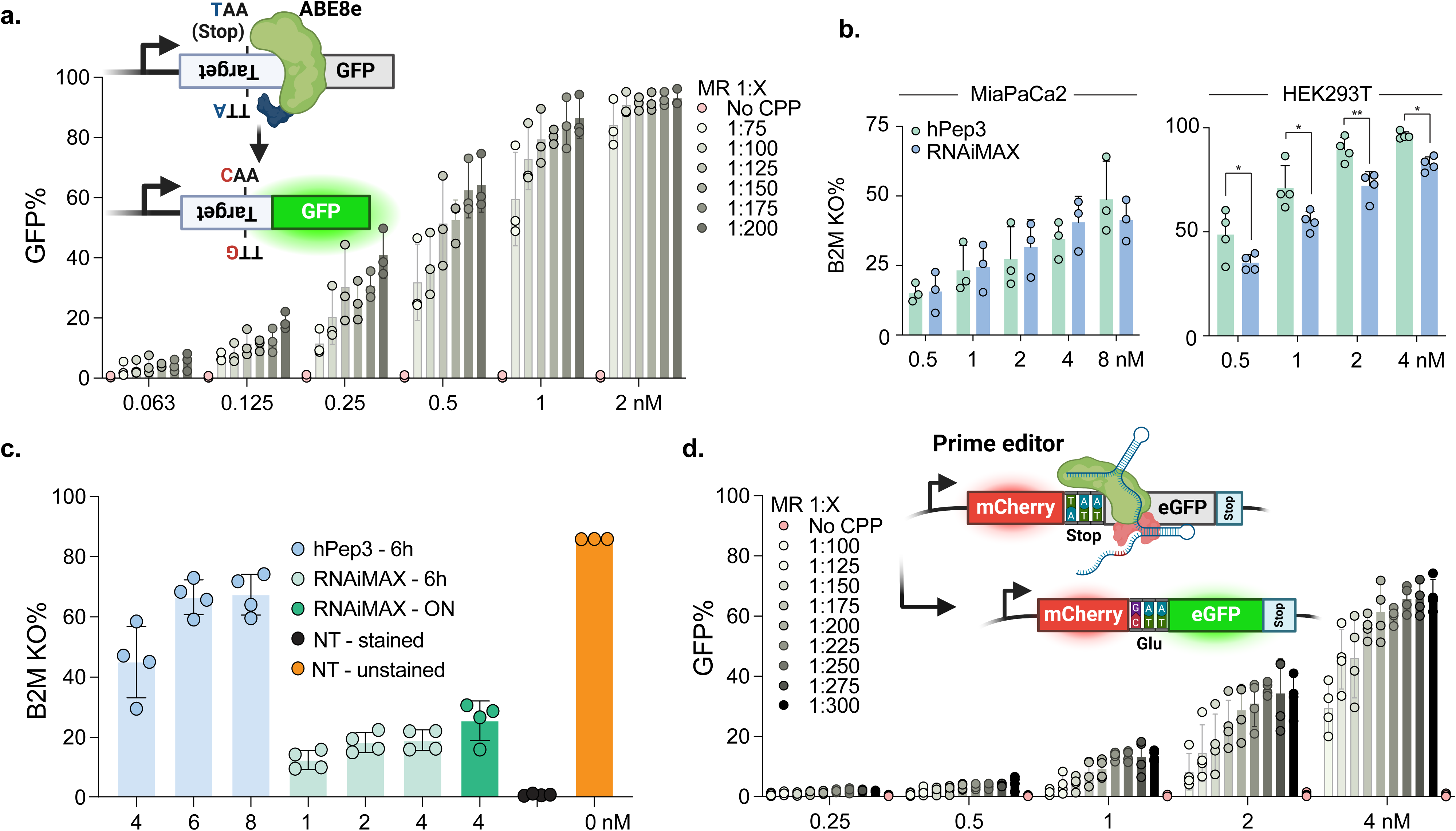

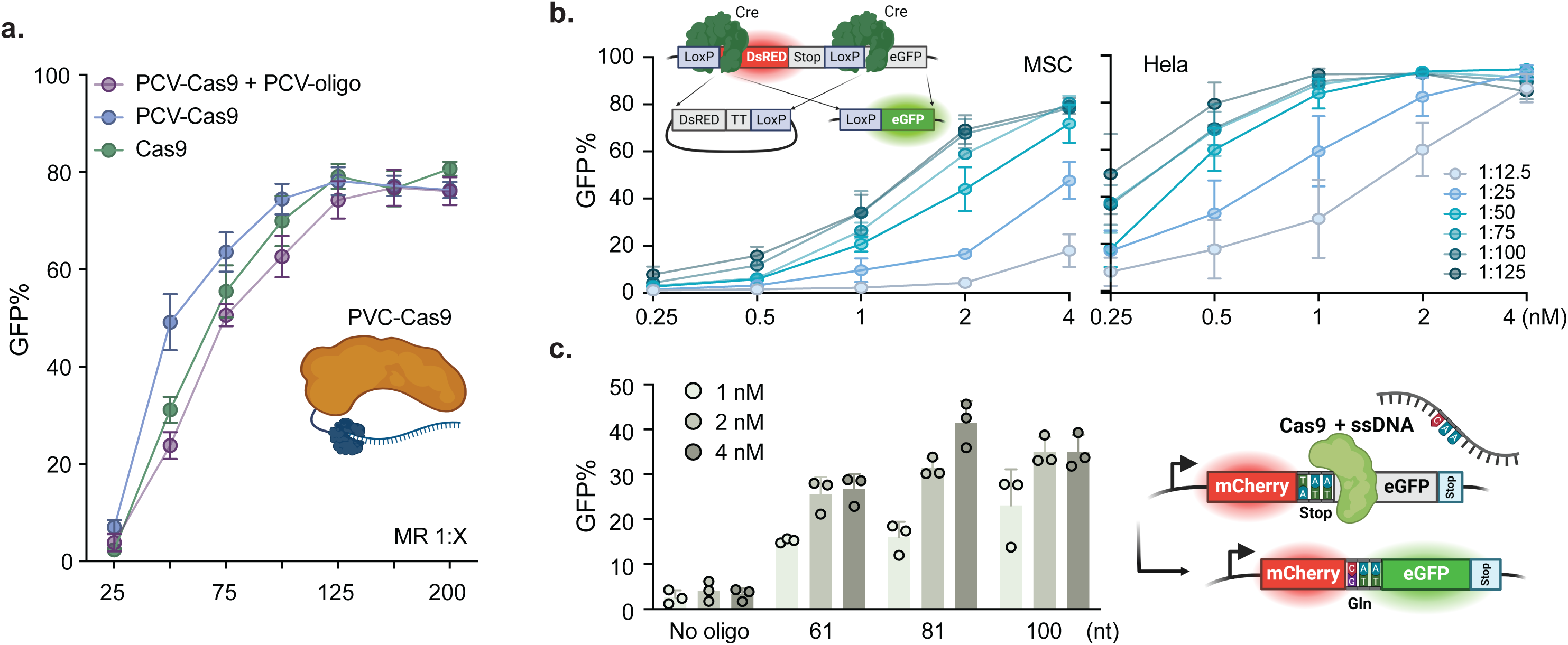

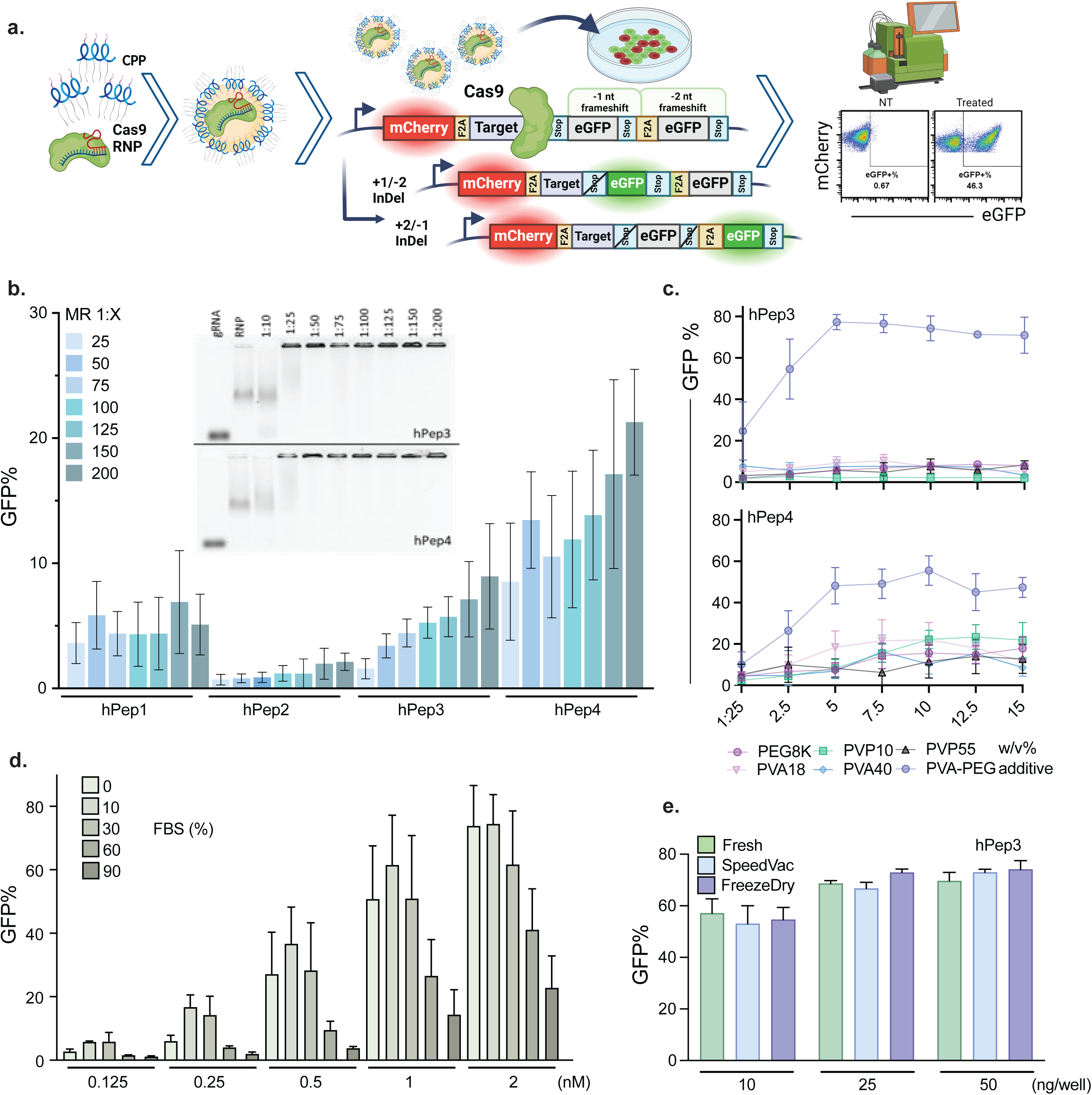

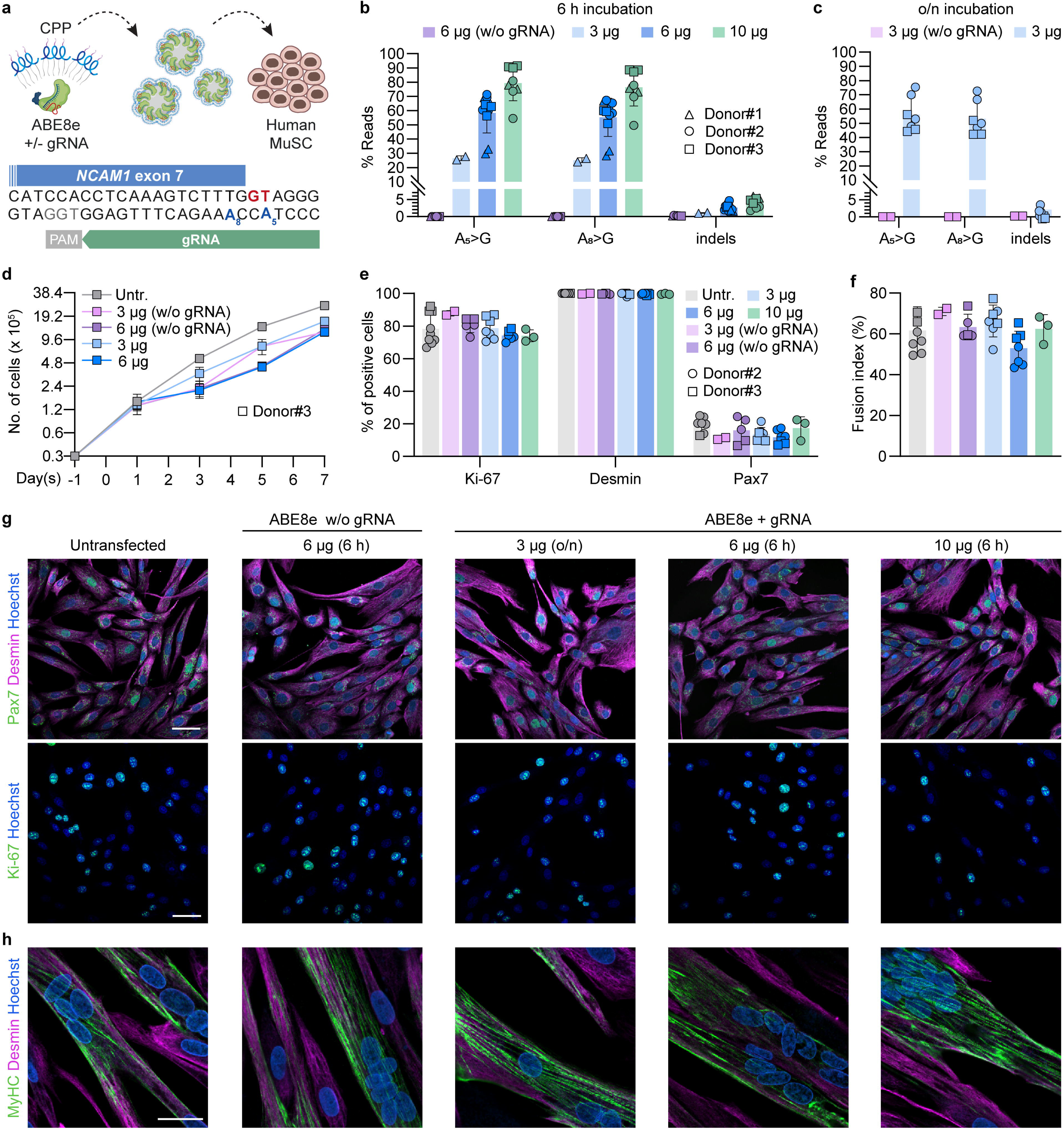

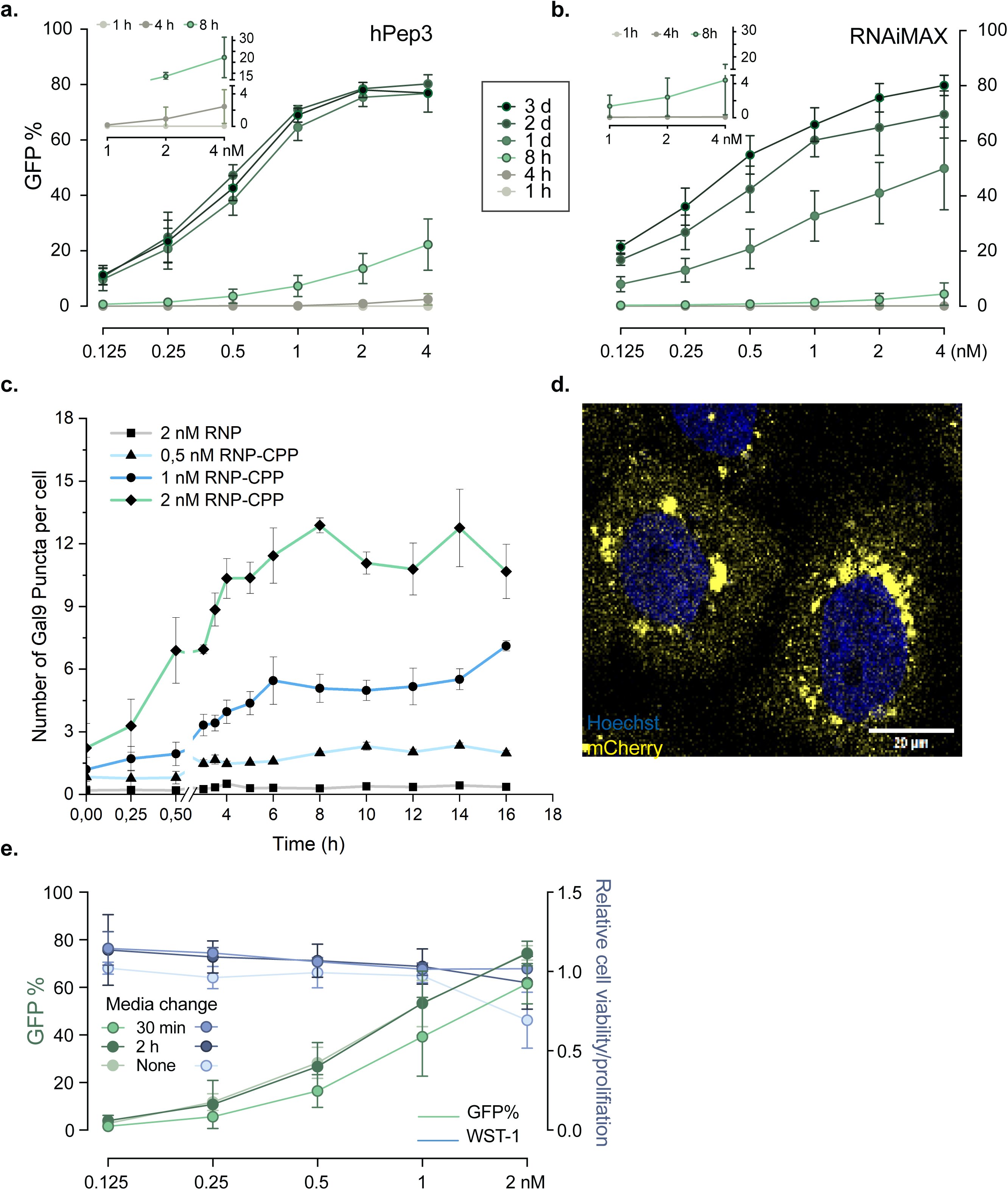

