## Supplemental figures for "Advanced Peptide Nanoparticles Enable Robust and Efficient delivery of gene editors across cell types"

**Supplementary information - Nanoparticle-based delivery of RNP gene editors using modified cell-penetrating peptides**

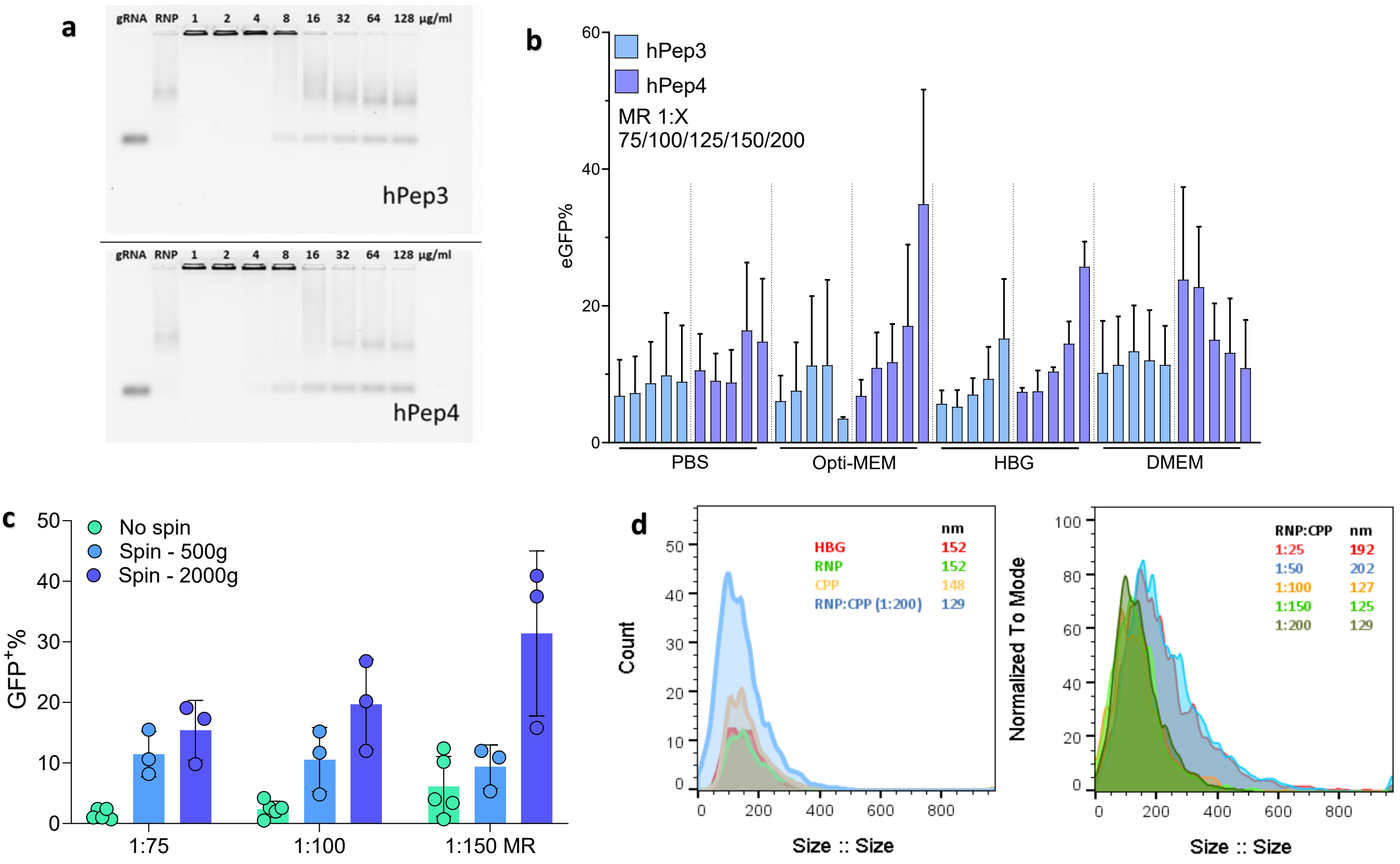


**Supplementary figure 1. Stability, size, and optimization of RNP:CPP complexes. a** Hexametaphosphate challenge of formed RNP-CPP particles (in HBG), where the hexametaphosphate disrupts both Cas9 – gRNA and RNP – CPP interactions. A larger amount of hexametaphosphate before complex disruption indicates more stable particles. **b** Testing of hPep3 and hPep4 in different buffers with increasing MR (200 ng Cas9 per 96-well. HEK293T SL cells). Mean value of n=3 independent experiments ± SD. **c** Spinfection testing of hPep:RNP particles (in HBG). The 96-well plates were spun for 30 min after adding nanoparticles (50 ng Cas9 per 96-well, HEK293T cells). Mean value of n=3 independent experiments ± SD. **d** ZetaView analysis of hPep3-Cas9 RNP particles formed in HBG. The left graph shows particle size and count. The right graph shows the normalized count and size of increasing RNP:CPP MR. At lower MR, the particles are more heterogeneous and larger, while at 1:100 and above, the population decreases in size down to around 130 nm.

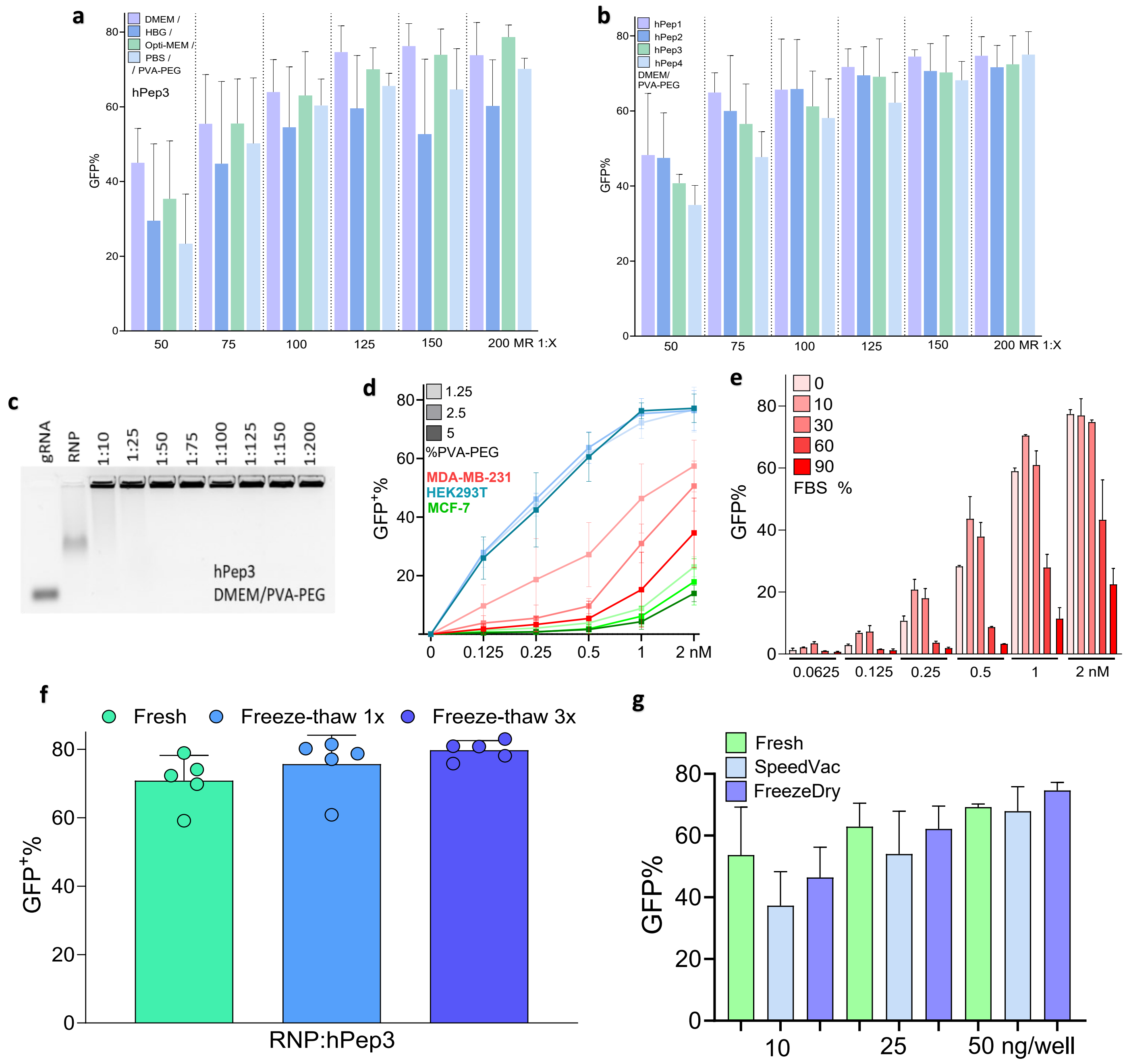


**Supplementary figure 2. Optimization and testing of Cas9:hPep3 in challenging conditions. a** Buffer testing when using PVA-PEG in the solution (5 w/v%) with increasing MR (25 ng Cas9 per 96-well, HEK293T cells). Mean value of n=3 independent experiments ± SD. **b** Testing of hPep1-4 in DMEM/PVA-PEG (5 w/v%) with increasing MR (25 ng Cas9 per 96-well, HEK293T cells). Mean value of n=3 independent experiments ± SD. **c** Gel mobility assay of RNP-hPep3 formed in DMEM/PVA-PEG. The RNP was formed using ATTO550-tagged trRNA. **d** RNP-hPep3 testing in challenging cell lines, MDA-MD-231 and MFC-7, with different w/v% of PVA-PEG in the solution. Mean value of n=3 independent experiments ± SD. **e** Testing of Cas9-hPep3 in increasing serum concentrations with a 2 h media change. Mean value of n=independent experiments ± SD. **f** Freeze-thaw testing of formed RNP-hPep3 complexes (50 ng per 96-well, HEK293T). Mean value of n=5 independent experiments ± SD. **g** Testing of RNP-hPep4 compatibility with different storage methods (10/25/50 ng Cas9 per 96-well, HEK293T cells). Mean value of n=3 independent experiments ± SD.

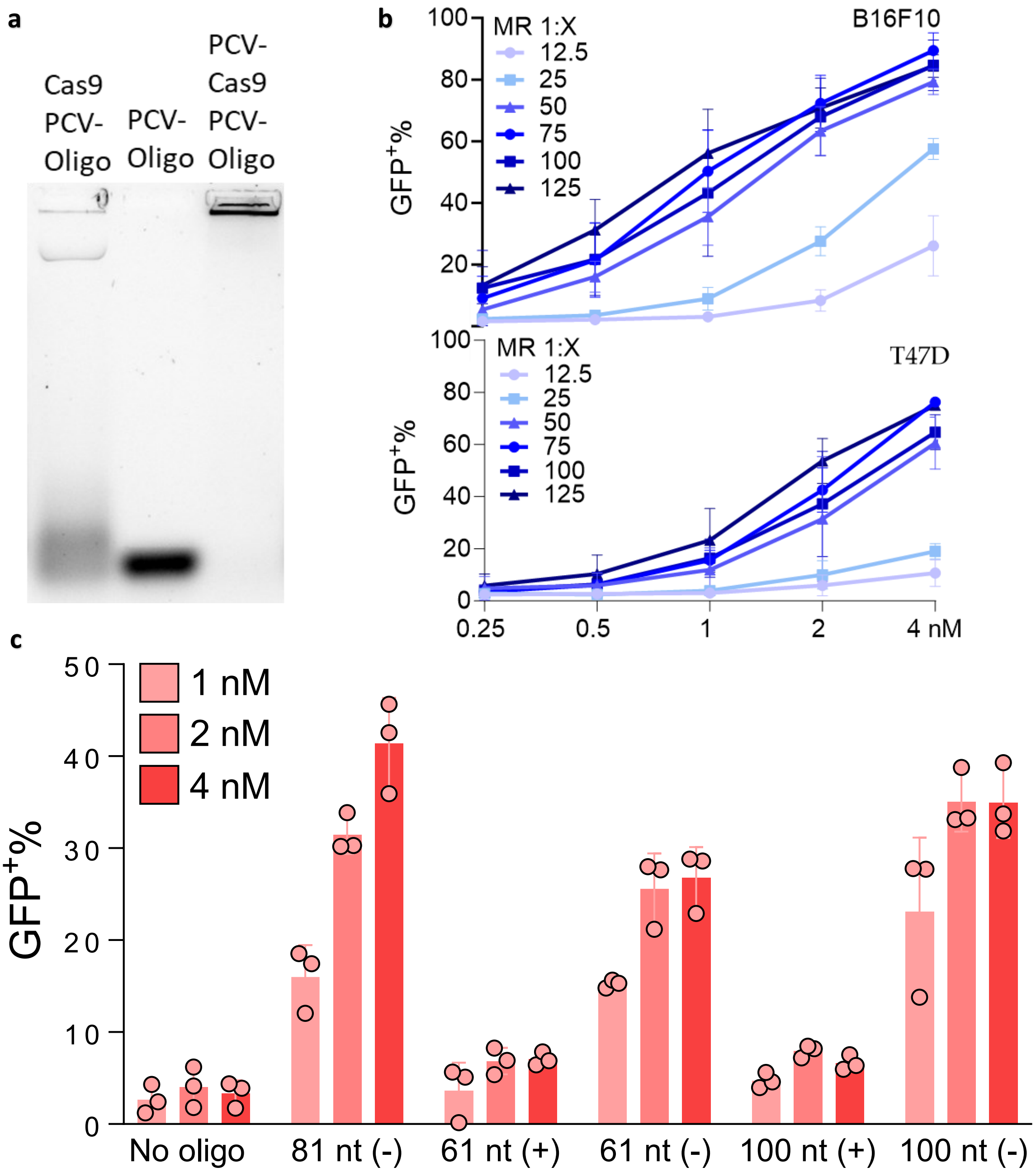


**Supplementary figure 3. Extended HDR results as well as Cre delivery. a** Gel electrophoresis of Cas9/PCV-Cas9 incubated with PCV-oligo showed that PCV-Cas9 is capable of binding to PCV-oligo. **b** Cre screening in breast cancer-derived B16F10 (murine) and T47D (human) cell lines harboring the TL reporter. Cre recombinase was complexed with hPep3 in DMEM/PVA-PEG buffer with increasing MR and added to cells in increasing dose. Mean value of n=3 independent experiments ± SD. **c** Cas9 mediated HDR screening as in **Fig2c** but now also displaying the ssDNA matching the (+) strand of the target DNA. With the editing rates achieved by the (+) binding oligos barely going over the no oligo control. Mean value of n=3 independent experiments ± SD.

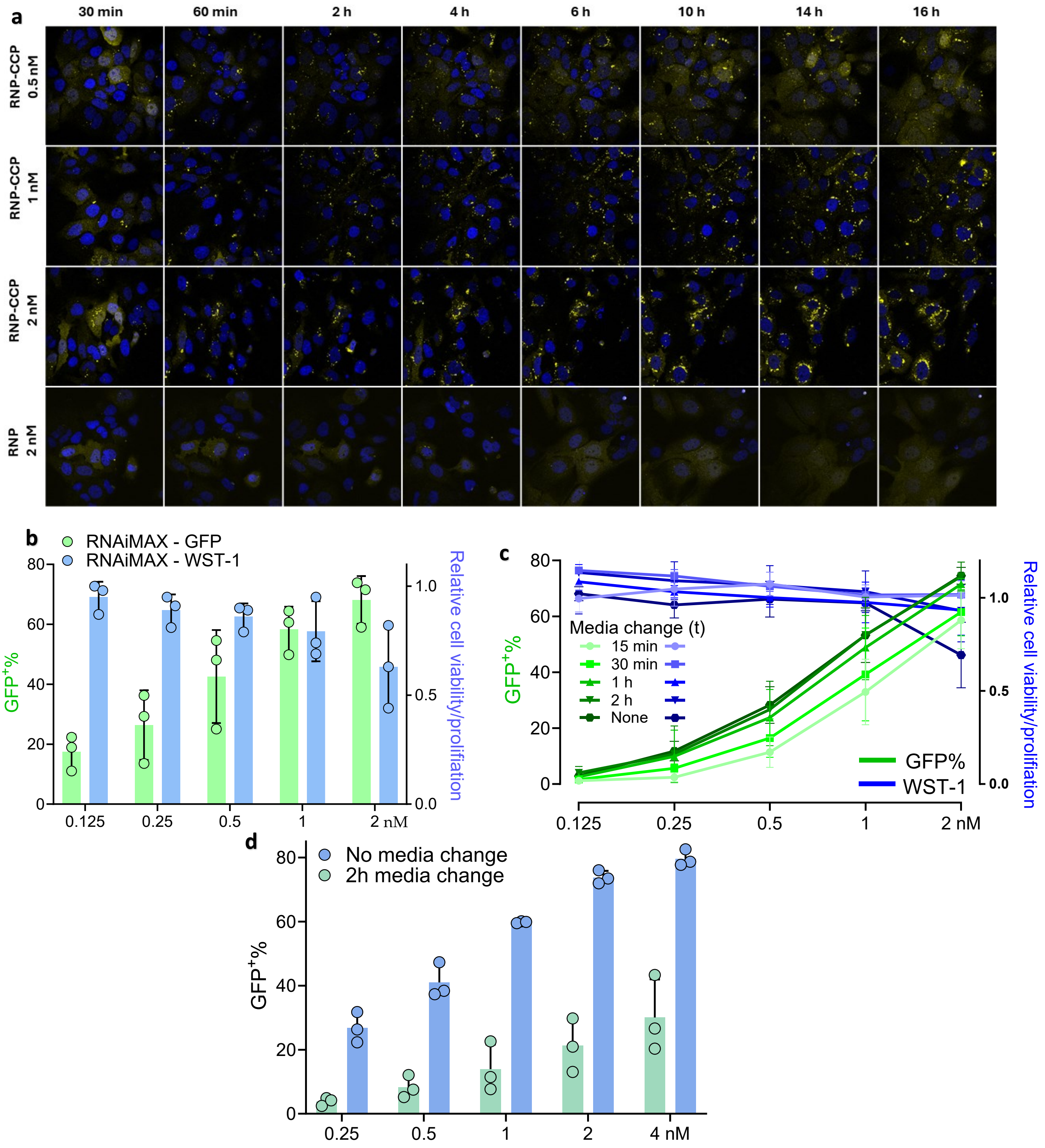


**Supplementary figure 4**. **Extended results showing the rapid endosomal rupture/gene editing and negation of adverse cellular effects**. **a** Representative images of Huh7 Gal9-mCherry reporter cells treated with RNP-hPep3 (MR 1:125) at increasing concentrations. Cells were imaged between 30 min and 16 h post-treatment, with puncta quantified at each time point. Hoechst (blue) and mCherry (yellow) were used. **b** RNAiMAX editing (green left y-axis) and WST-1 outcomes (blue right y-axis) with increasing doses added to HEK293T cells. Mean value of n=3 independent experiments ± SD. **c** Effect of medium change at different time points after RNP-CPP treatment of HEK293T SL cells on gene editing efficiency (green left y-axis) and WST-1 outcome (blue right y-axis). Mean value of n=3 independent experiments ± SD. **d** The kinetics of RNAiMAX transfection of HEK293T SL cells were tested by changing media at 2h. Media change for RNAiMAX strongly reduced transfection efficiency. Mean value of n=3 independent experiments ± SD.

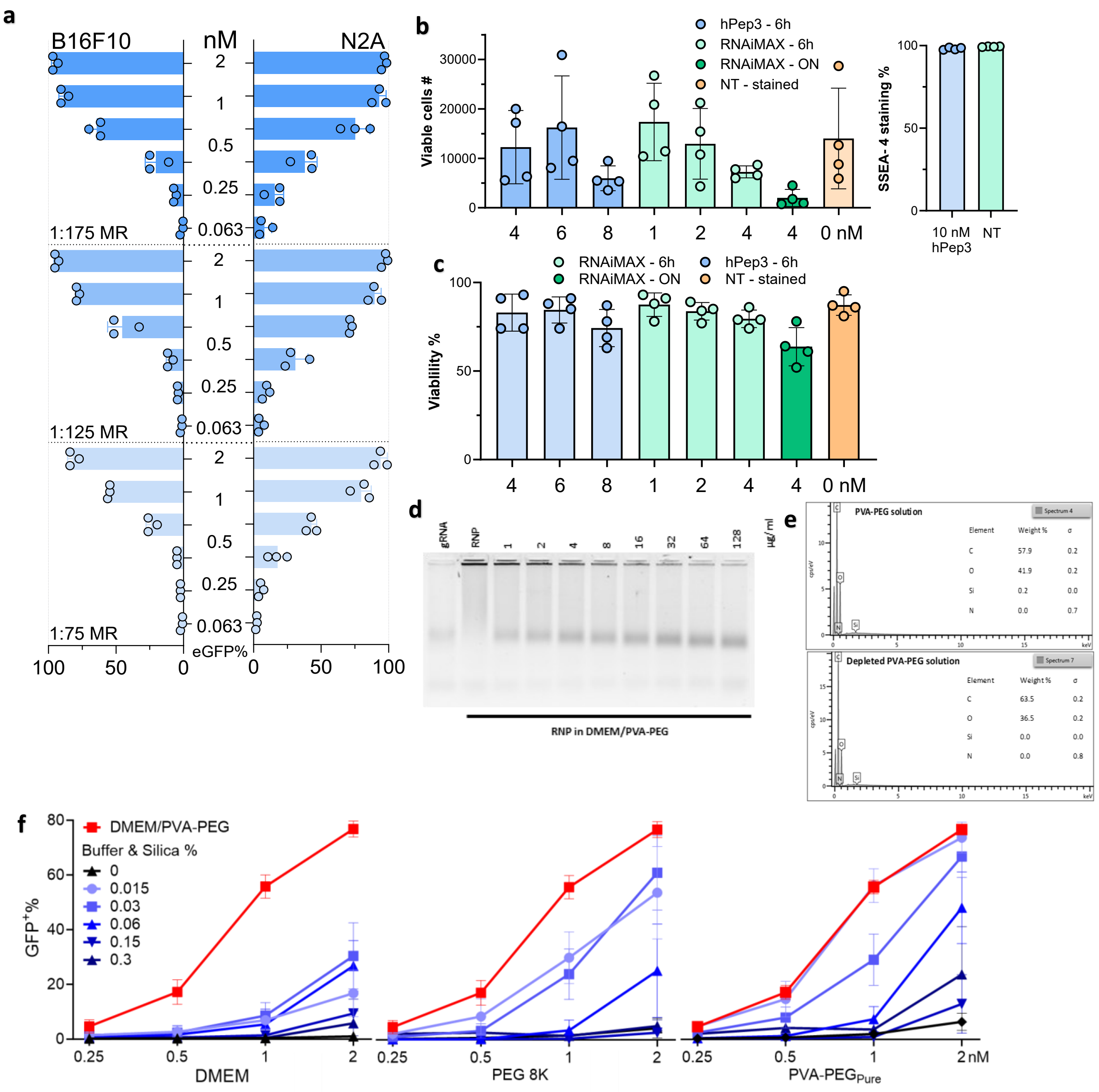


**Supplementary figure 5**. **Extended ABE8e-hPep3 delivery to B16F10/N2A/iPSC and characterization of PVA-PEG-associated silica**. **a** B16F10-ABE-GFP and N2A-ABE-GFP cells treated with ABE8e-hPep3 at increasing dose and MR. B16F10 and N2A reached 95%+ and 97%+ base conversion respectively. The highest editing reached in an N2A triplicate was 99.2%. Mean value of n=3 independent experiments ± SD. **b & c** Cell number and viability of iPSC cells treated with ABE8e:hPep3 or RNAiMAX as recorded by the flow cytometry. Results are from 4 different donor iPSC cells from 4 different donors done in triplicates, mean ± SD. Also shown is % of cells positive for the pluripotency marker SSEA-4, done with iPSC from 4 different donors in triplicates **d** Atto550 tagged Cas9 RNP was incubated in DMEM/PVA-PEG solution. The formed particles were then disrupted with increasing amounts of hexametaphosphate to investigate their stability. Disassociation was observed at the lowest tested concentration and increased with hexametaphosphate concentration. **e** Energy-dispersive x-ray spectroscopy investigated the elemental composition of the silica pellet and silica-depleted PVA-PEG. Images are representative of 3 replicates. Silica was not detected in any of the depleted samples. **f** PVA-PEG derived silica addition to different buffers and then used for RNP-hPep3 formulation followed by addition to HEK293T SL cells. With 0.015 w/v% matching the original concentration of silica in the PVA-PEG buffer. Mean value of n=3 independent experiments ± SD.

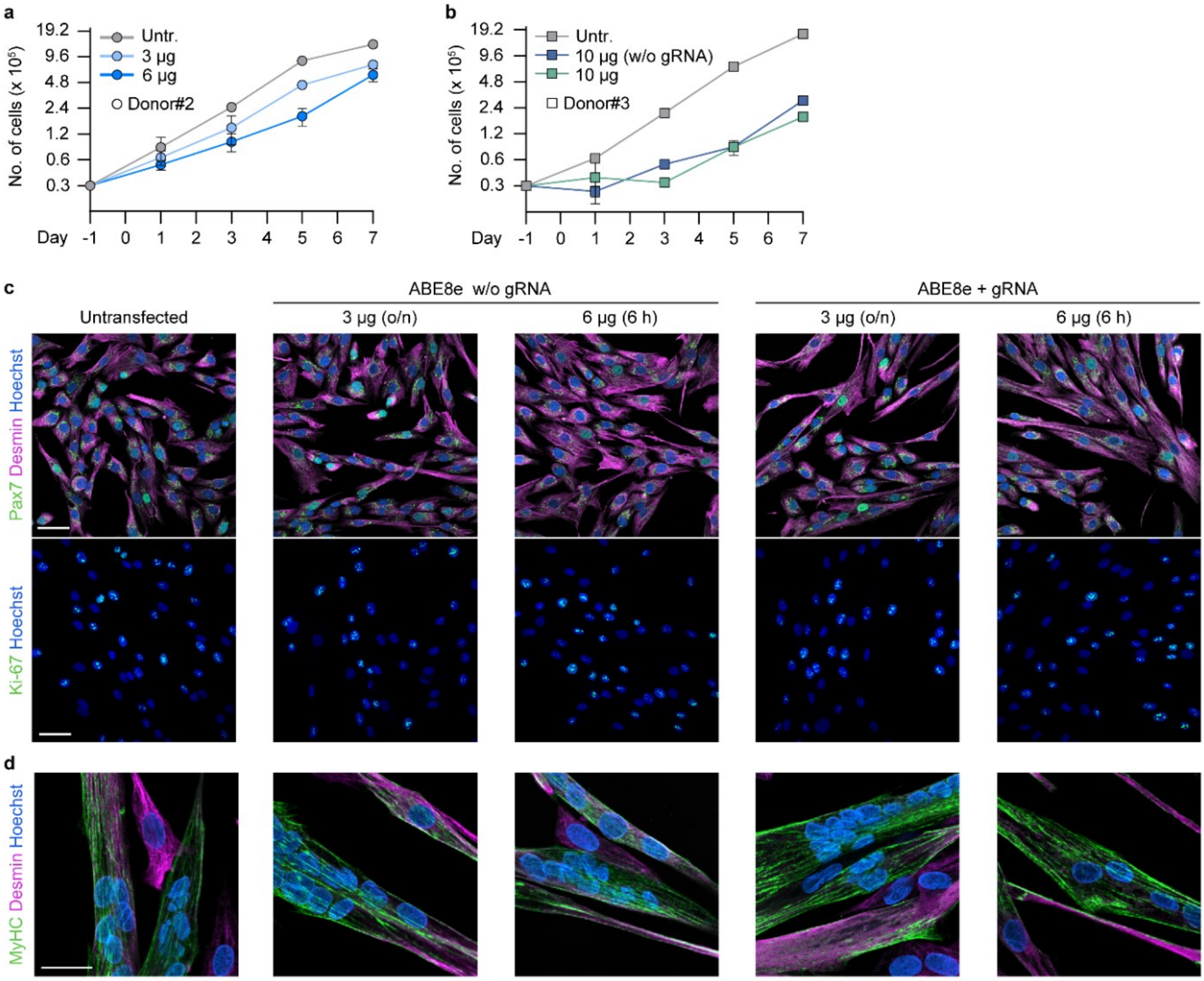


**Supplementary figure 6. MuSC growth and marker expression after treatment. a & b** Growth analysis of human MuSC over 7 days after treatment with hPep3-ABE8e targeting *NCAM1* exon 7 (*n* = 3 technical repeats per donor, mean ± SD). Cells were seeded on day -1 and treated at day 0. **c** Confocal microscopy images of MuSC from donor #3 immunostained for myogenic and proliferation markers after treatment with hPep3-ABE8e targeting *NCAM1* exon 7. Scale bars: 50 µm. **d** Confocal microscopy images of MuSC from donor #3 induced to fuse into multinucleated myotubes after treatment with hPep3-ABE8e. Scale bar: 50 µm. Untr.: Untransfected.

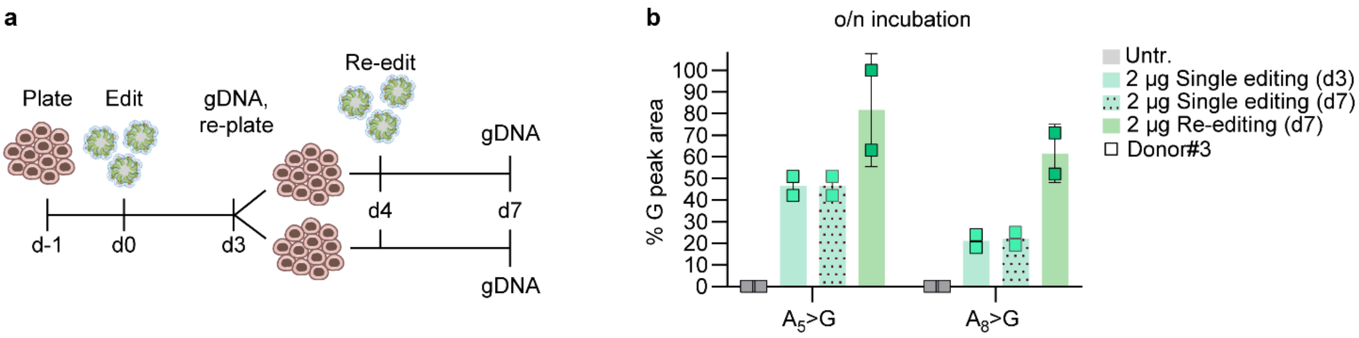


**Supplementary figure 8. Increasing editing in human MuSC with a second round of treatment of ABE8e:hPep3. a** Schematics of the experimental setup. **b** A>G conversion rates at adenines $A_5$ and $A_8$ in human MUSC treated with a single round (at day 0) or two rounds (day 0 + re-editing at day 4) of hPep3-ABE8e targeting *NCAM1* exon 7, determined by Sanger sequencing and EditR analysis (*n* = 2 technical repeats, mean independent experiments ± SD).
